# Human microRNA-153-3p targets specific neuronal genes and is associated with the risk of Alzheimer’s disease

**DOI:** 10.1101/2024.09.07.611728

**Authors:** Ruizhi Wang, Bryan Maloney, Kwangsik Nho, John S. Beck, Scott E. Counts, Debomoy K. Lahiri

## Abstract

Alzheimer’s disease (AD) is a progressive degenerative disease characterized by a significant loss of neurons and synapses in cognitive brain regions and is the leading cause of dementia worldwide. AD pathology comprises extracellular amyloid plaques and intracellular neurofibrillary tangles. However, the triggers of this pathology are still poorly understood. Repressor element 1-silencing transcription/neuron-restrictive silencer factor (REST/NRSF), a transcription repressor of neuronal genes, is dysregulated during AD pathogenesis. How REST is dysregulated is still poorly understood, especially at the post-transcriptional level. MicroRNAs (miRNAs), a group of short non-coding RNAs, typically regulate protein expression by interacting with target mRNA transcript 3’-untranslated region (UTR) and play essential roles in AD pathogenesis. Herein, we demonstrate that miR-153-3p reduces REST 3’-UTR activities, mRNA, and protein levels in human cell lines, along with downregulating amyloid-β precursor protein (APP) and α-synuclein (SNCA). We determine by mutational analyses that miR-153-3p interacts with specific targets via the seed sequence present within the respective mRNA 3’UTR. We show that miR-153-3p treatment alters the expression of these specific proteins in human neuronally differentiated cell lines and human induced pluripotent stem cells and that miR-153-3p is itself dysregulated in AD. We further find that single nucleotide polymorphisms (SNPs) within 5kb of the *MIR153-1* and *MIR153-2* genes are associated with AD-related endophenotypes. Elevation of miR-153-3p is associated with a reduced probability of AD, while elevated REST may associate with a greater probability of AD. Our work suggests that a supplement of miR-153-3p would reduce levels of toxic protein aggregates by reduced expression of APP, SNCA and REST expression, all pointing towards a therapeutic and biomarker potential of miR-153-3p in AD and related dementias.

## Introduction

Alzheimer’s disease (AD) and Parkinson’s disease (PD) are the two most common neurodegenerative diseases with distinct pathologies [1]. AD pathogenesis is characterized by the accrual of extracellular amyloid plaques composed mainly of amyloid-β (Aβ) peptides and intracellular neurofibrillary tangles composed of hyperphosphorylated tau protein in cognitive brain regions (e.g. hippocampus and cortex) [2]. By contrast, PD pathogenesis is characterized by Lewy bodies, which are mainly composed of α-synuclein, in the substantia nigra [3]. The occurrence of comorbid AD and PD pathologies resulting in Lewy body dementia is also common, as is the appearance of mixed pathologies in late-stage AD or PD [4, 5]. However, the triggers of these pathologies [6], especially for sporadic cases, are still poorly understood, and the identification of these triggers and their mechanism of action would promote the development of therapeutic interventions to slow neurodegeneration and synaptic loss and restore function. Although the *APOE* ε4 allele, which promotes Aβ aggregation and deposition, remains the major genetic risk factor for sporadic AD, growing evidence has shown that epigenetic regulation by microRNAs (miRNAs) plays a key role in AD pathogenesis [7–9]. Herein, we focus on molecular approaches to target specific genes involved in AD, reduce their potentially toxic protein products, and maintain neuronal health. One of the molecules tested is microRNA-153-3p (miR-153-3p), which may provide neuroprotection via its regulation of Repressor Element 1-Silencing Transcription (REST), also known as Neuron-Restrictive Silencer Factor (NRSF).

REST is a transcription repressor, mainly repressing neuron-specific gene expression during embryogenesis, neurogenesis and neuronal differentiation [10, 11]. REST represses target genes transcription by binding to their promotors, which include the type II sodium channel, SCG10, choline acetyltransferase, m4 muscarinic acetylcholine receptor, synapsin-1, and NgCAM [12–16]. Once REST binds to its target genes, it serves as a scaffold protein to recruit other repressor proteins such as Sin3, class I histone deacetylase (HDAC1 and HDAC2), methyl CpG binding protein 2 (MeCP2) and REST corepressor 1[17–21]. In mouse embryonic stem cells, REST helps maintaining self-renewal and pluripotency [22], whereas knockdown of REST in embryonic stem cells drives cell differentiation into the neuronal lineage [23]. Furthermore, activation of REST target genes by a recombinant dominant negative mutation REST-VP16 is sufficient to differentiate neural stem cells into functional neurons [24].

Recent studies found that REST also plays an important role in aging and AD. REST not only represses pro-apoptotic and AD-related genes but also provides neurons with resistance to Aβ and oxidative stresses.[25, 26] Reduced plasma levels of REST can serve as a potential biomarker for AD.[27] However, other studies found that REST levels are elevated in the brains of AD patients and associated with decreased level of Choline Acetyltransferase (ChAT) [28]. In cerebellum of AD mouse model, REST was increased together with reduced ChAT compared to those of wild-type animals [29].

Given the importance of REST, understanding how REST is regulated physiologically and dysregulated during aging and AD is critical. MicroRNAs (miRNAs) are a class of short non-coding RNAs with length around 22 nucleotides [30]. Notably, miRNAs generally downregulate target mRNAs expression by base paring with their 3’ untranslated regions (UTR) with imperfect complementarity and recruiting RNA-induced silencing complex (RISC) to either degrade target mRNAs or inhibit target mRNAs translation [31]. miRNAs contribute to or protect against neurodegenerative disease progression by modulating key pathological or beneficial proteins [32]. For example, the Aβ precursor protein (APP) is extensively regulated by miRNAs, including miR-20b, miR-101, miR-31, miR-346, miR-106a, and miR-520c [33–40], while β-site amyloid precursor protein cleaving enzyme 1 (BACE1) is also regulated by many miRNAs including miR-124, miR-339, miR-298 and miR-29c [41–45]. These results underscore the potential clinical efficacy for microRNA-based therapies for AD and related dementias (ADRD) [46–48].

In this report we used four major approaches: 1) Bioinformatics tools, 2) Cellular alterations of target mRNAs/proteins via miR-153-3p transfection in cell cultures, 3) Disease-specific alterations in miR-153-3p in brain tissue specimens from control and AD subjects, and 4) Genomics and association studies. We show that 1) miR-153-3p interacts with specific targets via the seed sequence present within their respective mRNA 3’UTRs, 2) miR-153-3p treatment alters the miR-153-3p network proteins in human induced pluripotent stem cells (hiPSCs), 3) miR-153-3p is dysregulated in AD; and 4) SNPs near *MIR153* genes associate with AD endophenotypes.

Bioinformatics prediction tools reveal multiple miRNAs that target the REST mRNA 3’-UTR. Among these miRNAs is hsa-miR-153-3p (miR-153-3p). This miRNA reduces levels of APP in cell cultures [36] and is itself dysregulated in AD patient brains as well as in the APPswe/PSΔE9 mouse model of AD [35, 36]. Further connections between miR-153-3p and AD include the presence of an AD-associated single nucleotide polymorphism (SNP) in the APP mRNA 3’-UTR that crosses an miR-153-3p binding site [49, 50] and an SNP in the miR-153-3p site that may be protective vs. AD [50]. Beyond its effects on AD, miR-153-3p is also involved in PD pathology. It reduces α-synuclein expression and protect neurons against neurotoxin 1-Methyl-4-Phenyl-Pyridinium (MPP+) through mTOR pathway [51, 52], and miR-153-3p is elevated in cerebrospinal fluid (CSF) of PD patients [53]. Salivary and plasma miR-153-3p levels can serve as potential biomarkers of PD [54, 55]. Moreover, miR-153-3p improves cognitive outcomes following ischemic stroke [56]. Functionally, miR-153-3p is also associated with synaptic vesicular transport and release as well as short-term memory deficit [57–60]. Mature miR-153-3p is cleaved from products of two genes, specifically *MIR153-1* on chromosome 2 and *MIR153-2* on chromosome 7. *MIR153-1* appears to only produce miR-153-3p from its transcribed hairpin [61], while *MIR153-2* produces both miR-153-3p and miR-153-5p [62].

Herein, we report that elevation of miR-153-3p levels associated with greater risk of AD, specifically in temporal lobe and posterior cingulate cortex (PCC). We consulted multiple online bioinformatics prediction tools and found that miR-153-3p may target REST, APP and SNCA mRNA 3’-UTRs. We constructed reporter plasmids containing 3’-UTR sequences for these genes and determined that administering miR-153-3p reduced reporter activities for all three, in a time-dependent manner. Antagomirs to miR-153-3p reversed these effects and increased reporter activities. When we treated several human cell cultures, specifically epithelial (HeLa), neuroblastoma (SK-N-SH) cells that were neuronally differentiated (Diff-NB), glioblastoma (U373), microglial (HMC3), and neuronal stem cells which are derived from a normal human induced pluripotent stem cell line (iPSC), with miR-153-3p, it reduced native endogenous REST mRNA and protein levels. Moreover, miR-153-3p treatment induced neuronal differentiation in neuronal stem cells.

## Materials & Methods

### Human brain samples from AD and age-matched non-cognitively impaired (NCI) controls

Age-matched samples of temporal lobe (BA 21/22) and cerebellum from control (CTL) and AD subjects representing both sexes were obtained from the University of Kentucky AD Research Center Brain Bank (n = 48), whereas posterior cingulate cortex (BA 23) was obtained from the Rush AD Research Center Brain Bank (n = 39). Exclusion criteria for cases selection at both sites included evidence of synucleinopathies such as Parkinson’s disease and Lewy body disease, frontotemporal dementia, argyrophilic grain disease, vascular dementia, hippocampal sclerosis, and/or large strokes or lacunes.

### MicroRNA quantification by qRT-PCR

RNA was extracted from frozen tissue using a modified Ambion PureLink mini kit protocol (#12183018A). Briefly, between 10 to 25 mg of tissue was placed in a 2 ml round bottom tube and 1 ml of Trizol (ThermoFisher #15596026) was added. Tissue was sonicated on ice until homogenous and was allowed to incubate for 5 minutes at room temperature. Then 200 μl of chloroform was added, and the sample was vortexed for 15 seconds. Following a 3-minute incubation at room temperature, the samples were centrifuged at 12000 x g for 15 minutes at 4° C. The upper aqueous layer was transferred to a clean 1.5 ml tube, and an equal volume of 70% ethanol was added. The sample was vortexed and then processed following the manufacturer’s instructions. RNA was eluted in a final volume of 50 μl of nuclease free water and was then quantified to be used as a template for cDNA synthesis.

### Quantitation of miR-153-3p levels

miR-153-3p levels in human tissue were analyzed by qPCR using both relative and absolute quantitative techniques. For relative quantitation, a probe-based assay for miR-153-3p (TaqMan 001191) was measured and compared to the control small RNA RNU48 (TaqMan 001006 labelled with VIC) [63]. Briefly, template for qPCR was generated using the TaqMan microRNA reverse transcription kit (Applied Biosystems 4366596) following the manufacturer’s recommended protocol with an input of 10 ng of RNA. qPCR was performed on an ABI 7500 instrument in 20 μl reactions, which were incubated for 40 amplification cycles. Each reaction contained 1.3 μl of reverse transcription product as template, 2x master mix minus UNG (Applied Biosystems 444040), and each of the TaqMan assays listed above. Ct values were determined using a constant threshold, and fold change was calculated by the delta-delta Ct method.

For absolute quantification, TaqMan Advanced cDNA synthesis kit (Applied Biosystems A28007) was used to produce template for qPCR. 10 ng of RNA was poly adenylated, ligated to an adapter, reverse transcribed, and amplified, resulting in cDNA capable of being interrogated with any TaqMan Advanced miR assay. qPCR amplification reactions were assembled including 2 μl of miR-AMP product as template, 2x PrimeTime master mix (Integrated DNA Technologies 1055772), and TaqMan Advanced assay miR-153-3p (Applied Biosystems 477922) in a total of 10 μl. The reactions were subjected to 40 rounds of amplification in an ABI 7500 thermocycler. A standard curve of not less than five data points was created using known concentrations of a miR-153-3p synthetic oligonucleotide (IDT). Ct values were determined using a constant threshold. Construction of the standard curve was performed by creating a scatter plot in Excel based on the Ct values of the samples of synthetic miR-153-3p. The x-axis of the plot was converted to log scale, and a logarithmic trendline was fitted to the standards. The equation of the slope and the R squared value was displayed on the graph. Concentrations of unknown samples were determined by extrapolation using the slope equation generated by the standard curve.

### Single-nucleotide polymorphisms in proximity to MIR153 gene in the Alzheimer’s Disease Neuroimaging Initiative (ADNI) cohort

The ADNI initial phase (ADNI-1) was launched in 2003 to test whether serial magnetic resonance imaging (MRI), position emission tomography (PET), other biological markers, and clinical and neuropsychological assessment could be combined to measure the progression of mild cognitive impairment (MCI) and early AD [64, 65]. ADNI-1 has been extended in subsequent phases (ADNI-GO, ADNI-2, ADNI-3, and ADNI-4) for follow-up of existing participants and additional new enrollments. More information about ADNI can be found at https://adni.loni.usc.edu. Demographic information, pre-processed neuroimaging scans, *APOE* and whole-genome genotyping data, neuropsychological test scores, and clinical information are publicly available from the ADNI LONI data repository. The ADNI participants were genotyped using several genotyping platforms (Illumina Human610-Quad BeadChip, Illumina HumanOmni Express BeadChip, Illumina Infinium Global Screening Array BeadChip, and Illumina HumanOmni 2.5M BeadChip) [66, 67]. Un-genotyped SNPs were imputed separately in each platform using the Haplotype Reference Consortium (HRC) data as a reference panel. Before imputation, standard sample and SNP quality control (QC) procedures were performed: (1) for SNP, SNP call rate < 95%, Hardy-Weinberg P value < 1×10^−6^, and minor allele frequency (MAF) < 1%; (2) for sample, sex inconsistencies, and sample call rate < 95% [66, 68]. In order to prevent spurious association due to population stratification, only non-Hispanic participants of European ancestry by multidimensional scaling analysis were selected, which clustered with HapMap CEU (Utah residents with Northern and Western European ancestry from the CEPH collection) or TSI (Toscani in Italia) populations using multidimensional scaling (MDS) analysis (www.hapmap.org) [69]. Imputation and QC procedures were performed as described previously [68]. We then consulted the SNP database “dbSNP”, maintained by the National Library of Medicine, USA [70] to confirm publication of SNPs in the population at large.

### Association of SNPs with selected AD-associated endophenotypes in the ADNI cohort

In addition to genotyping, the following endophenotypes were investigated in the ADNI cohort. Cognitive scores, specifically composite scores for memory [71, 72] the Alzheimer’s disease assessment scale-13 item (ADAS13) [73], mini-mental state exam (MMSE) [74], and the Rey auditory verbal learning test (RAVLT) [75]; cerebrospinal fluid (CSF) levels of tau and Aβ measured by using the validated and highly automated Roche Elecsys® electrochemiluminescence immunoassays; brain deposition of Aβ plaque, a global cortical amyloid standardized uptake value ratio (SUVR) normalized to whole cerebellum measured by Amyloid PET Brain glucose metabolism, a global cortical glucose metabolism SUVR normalized to pons, measured by FDG-PET [76]; and entorhinal cortical thickness and hippocampal volume measured by MRI [76]. Thes cognitive, biomarker, and anatomical measurements were tested as response variables to presence of specific SNPs detected by SNPs within 5kb up or downstream of the *MIR153-1* and *MIR153-2* genes.

### Identification of putative miR-153-3p binding sites on the REST and other mRNA 3’UTRs

Multiple bioinformatic prediction tools with different algorithms, specifically TargetScan [77], DIANA-microT [77], miRDB [78], RNA22 [79], StarMir [80, 81], and miRmap [82] were applied to find the binding site of miR-153-3p in REST mRNA 3’UTR.

### Cell cultures

Human glioblastoma cells (U373), HeLa cells, neuroblastoma cells (SK-N-SH), and microglia cells (HMC3) were obtained from ATCC. Neuroblastoma cells were further treated with retinoic acid to produce differentiated neuronal cells (Diff-NB) or neuronotypic cells [83]. Cells were grown in Eagle’s modified minimum essential media (EMEM) containing 10% FBS and penicillin/streptomycin solution at 37 °C in 5% CO2 humid incubators.

### Plasmid and RNA Transfection

Transfections were performed when cells reached around 80% confluency. Culture media were replaced with Opti-MEM media with 1% FBS and antibiotics were omitted from transfection media. For miRNAs and siRNAs transfection, Lipofectamine RNAiMax was applied 2ul per well in a 24 well plate format. 75nM miRNA or 50nM siRNA were premixed with RNAiMax according to its protocol. Mixes were added into each well and kept for 72 hours or otherwise indicated in figure legends. For co-transfection of plasmid with miRNA, plasmids were added at 100ng/well in a 96-well plate or 400ng/well in 24 plates. RNAiMax and miRNA were scaled down or up accordingly. Luciferase assay plasmids were obtained from Genecopia. MiR-153 mimics and antagomiRs were purchased from Dharmacon, Inc, mature miR-153 sequence (UUGCAUAGUCACAAAAGUGAUC).

### Cell lysing and content harvest

After washing with PBS, cells were lyses on-plate with vigorous shaking using 100ul RIPA buffer containing 1X halt protease inhibitors cocktail (Thermo Scientific #78430) except for extraction of α-synuclein, for which 8M urea buffer containing 0.5% SDS and protease inhibitors was used. Protein concentration was determined by BCA assay according to their protocol and then Laemmli sample buffer (LSB) was added to each tube of lysate. Lysate and LSB mixes were boiled for 10 minutes and cooled down on ice or kept in freezers for further studies. For proteomics study, cell pellets were collected and immediately frozen in liquid nitrogen. Then urea buffer was applied to lyse the cells and extract proteins.

### SDS–polyacrylamide gel electrophoresis (SDS–PAGE) and western blotting

An equal amount of protein lysate was loaded onto 26 lane Bis-Tris XT denaturing 4-12% polyacrylamide Gels and run with XT MOPS or XT MES buffer. Proteins were separated with SDS-PAGE and then transferred overnight onto PVDF membranes. Membranes were stained with 0.1% Ponceau S solution to confirm transfer success. After three times of washing with TBST, membranes were incubated with 5% nonfat milk in TBST for 1 hour at room temperature. Primary antibodies were incubated at either room temperature for 3 hours or 4 °C overnight. Goat anti rabbit or mouse secondary antibodies were applied for 1 hour at room temperature. Protein bands were visualized using auto radiographic films and ECL buffer. Films were scanned for densitometry analysis. Antibodies used in this study: REST (Millipore 07-579), APP (Millipore MAB348), SNCA (ThermoFisher AHB0261), Doublecortin (Cell Signaling 4604), Nestin (Rockland 600-401-417), neuron-specific enolase (NSE) (Abcam ab16873) and β-actin (Millipore A5441).

### RNA-sequencing analysis

mRNA samples were extracted from ATRA differentiated neuroblastoma cells SK-N-SH (Diff-NB) with mirVana miRNA isolation kit. RNA quality was confirmed with a threshold of RNA integrity number (RIN) above 7 and 28S/18S rRNA ratio above 1. RNA samples were shipped to BGI Americas for RNA-seq, in which DNBSEQ transcriptome profiling was applied.

### Human Induced Pluripotent Stem Cells (iPSCs) culture

Human iPSC cells were obtained from Coriell Institute for Medical Research and maintained with mTeSR plus (STEMCELL Technologies). iPSC cells were subsequently differentiated into neural progenitor cells (NPCs), forebrain neurons and astrocytes with specific induction media and kit according to manufacturer’s instructions. Protein and mRNA were harvested at different stages for further analysis. Additional neural stem cells were obtained from Applied StemCell, Inc and maintained in NSC maintenance media (Applied StemCell, Inc) according to the manufacturer’s instructions.

### Proteomic screening of differentiated neuroblastoma (Diff-NB) cells treated with miR-153-3p

Lysates of cells treated with miR-153-3p, with antagomiR to miR-153-3p, and with mock transfection were subjected to Tandem mass tags (TMT) global proteomics analysis. 30 μg of each sample was reduced, alkylated, digested and labeled with 0.2 mg of TMT. High pH basic fractionated into 9 fractions and run on Lumos Orbitrap. Proteins with abundances that were reduced vs. mock by miR-153-3p and increased vs. mock by antagomiR or those with abundances increased vs. mock by miR-153-3p and reduced by antagomiR were selected fur further screening. Of these outputs, proteins with a relative log base 2 abundance of treatment vs mock less than 0.2 or p value less than 0.05 were further excluded. The remaining protein gene IDs were included alongside APP, DCX, NES, REST, and SNCA for network analysis.

### Network analysis of proteomic outputs

Filtered proteomic output identifications were entered into the NetworkAnalyst utility and database [84] and protein-protein interaction networks were constructed for hippocampus and frontal cortex. Steiner Forest networks were selected to produce more focused networks. After network creation, nodes were loaded into String DB [85] to evaluate strengths of node links and determine gene ontologies, network STRING clusters, KEGG and reactome pathways, and human disease phenotypes.

### Database search of miRNA levels in human tissue

To compare our measurements of miR-153-3p levels in various cell lines to human tissue, we consulted miRNATissueAtlas2 [86] for levels of miR-153-3p in various human tissues. We excluded any tissues that did not have at least three measurements and performed explicit comparisons.

### Data Analysis

All analyses were done with the R environment [87] and the arm [88], lme4 [89], MuMIn [90], emmeans [91], cluster [92] and Hardy-Weinberg [93] packages. Relationships between brain mRNA/miRNA levels and NCI/AD diagnosis was analyzed by logistic regression of miR-153-3p and REST RNA levels vs. NCI/AD diagnosis. We also tested potential covariates such as subject age, sex, and *APOE* genotype score. *APOE* genotype score was generated by simply scoring potential risk for each allele in a subject’s *APOE* genotype according to relative risks from population studies [94]. These scores corresponded one-to-one to specific genotypes. Models were compared via the second-order Akaike information criterion (AICc) [95]. We tested allele equilibrium for SNPs with the Hardy-Weinberg test adapted for stratification by sex and presence of at least *APOE*ε4 allele [96]. Each SNP proximal to the *MIR153-1* and *MIR153-2* genes were compared to the endophenotypes along with the potential covariates of subject sex, age, formal education (years), and presence of at least one *APOE*ε4 allele. Models were compared via AICc, selecting the most parsimonious model for each endophenotype for statistical testing and interpretation.

Luciferase reporter assays were analyzed by generalized linear models (glm) if endpoint or by segmented generalized linear mixed-level models (glmer) if time course experiments. Based on visual expression of data, time courses were divided into three segments that we named “Acute Response” (R), “Lag” (L), and “Continuation” (C). Western blots and CTG were analyzed by glm unless multiple blots were combined, at which point glmer were used to account for blot-to-blot variation. In all cases, olr, glm, and glmer were followed by ANOVA. When appropriate, pairwise comparisons were performed by estimated marginal means (emm) or estimated marginal trends (emt), using the Benjamini-Hochberg false discovery rate adjustment.

Proteomic abundance differences were filtered on two axes: Differences with a value between −0.2 and 0.2 and base 10 logarithm ≥ 1.303 were selected for further analysis. The resulting protein IDs were combined with *APP*, *DCX* (doublecortin), *NES*, *REST*, and *SNCA*. The combined list was used as input obtain network analysis via the NetworkAnalyst utility [84].

## Results

### hsa-miR-153-3p levels in temporal lobe and PCC associate inversely with probability of AD

We measured miR-153-3p levels from human brain samples that included TL and PCC samples by qRT-PCR as described. miR-153-3p levels were then modeled with *REST* mRNA levels in the same regions from the same cases to assess their associations with AD risk. *APOE* genotype and AD risk were also assessed for comparison (Table 1, Fig. 1). In our model, the probability of AD decreased as miR-153-3p levels increased. On the other hand, the probability of AD was modestly increased as REST mRNA levels increased. Finally, the probability of AD increased as individual *APOE* genotype risk scores increased, as expected [97, 98].

**Table 1.** ANOVA of Effects of miR-153-3p and REST on AD risk.

| **Effect** | **$\chi^2$ (df)** | **p** | **Tjur's *D*** |
| --- | --- | --- | --- |
| miR-153-3p | 20.5 (1) | < 0.001 | 0.259 |
| *APOE* Risk | 5.7 (1) | 0.017 | 0.092 |
| Brain Region | 6.0 (1) | 0.014 | 0.078 |
| REST | 4.7 (1) | 0.030 | 0.058 |

**Fig. 1.**
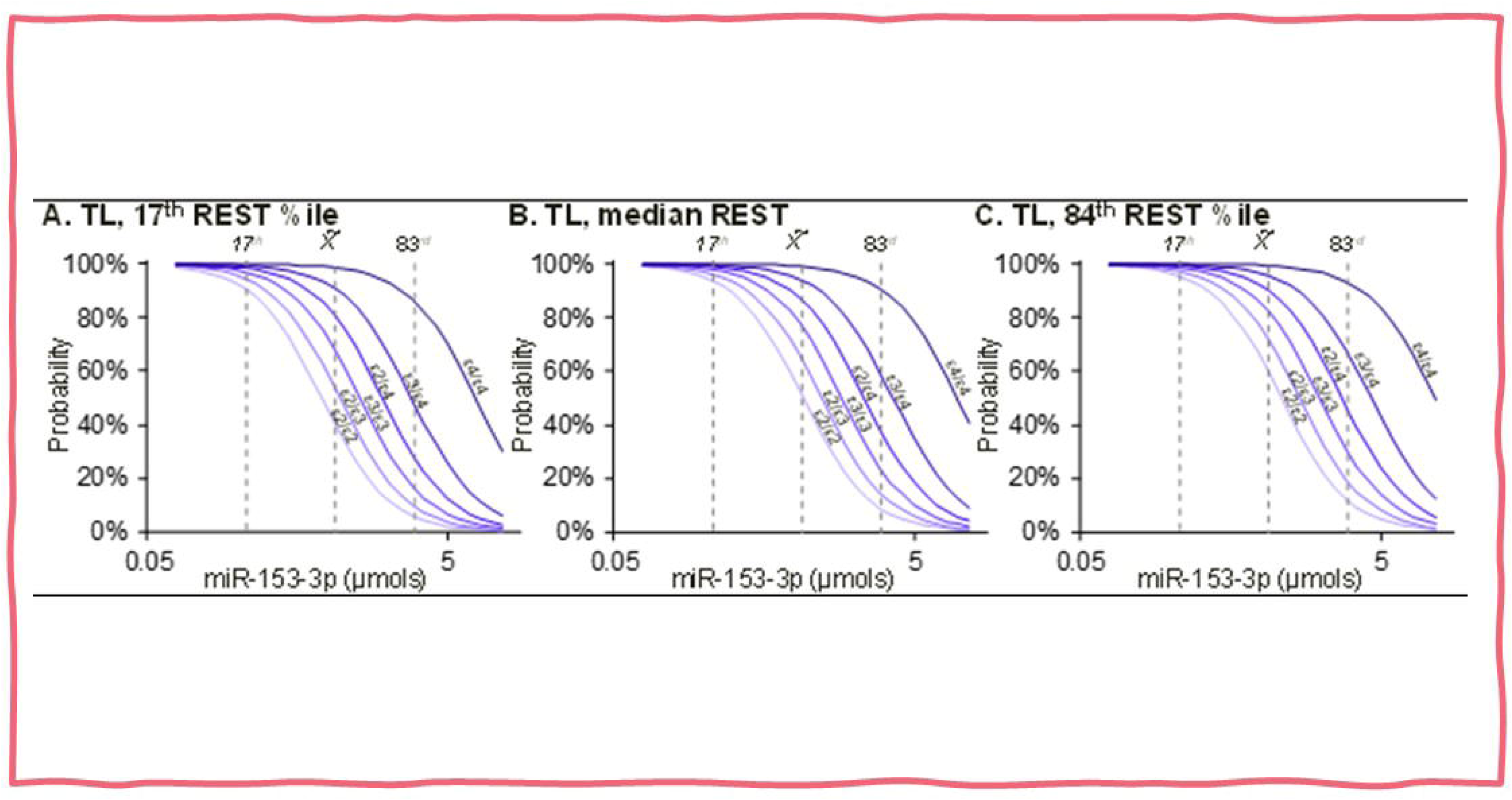
Probability of AD is reduced as levels of miR-153 increase, REST partially reverse this effect. RNA was extracted from human brain samples and subject to qRT-PCR as described in the text. A logistic regression model was constructed that also took into account effects of *APOE* genotype and brain region. Models revealed that each variable exerted significant effect on probability of AD. *APOE* genotype resulted in greater probability of AD as genotype associated risk increased from ε2/ε3 → ε3/ε3 → ε3/ε4 → ε4/ε4. These were used to predict estimated AD risk for other genotypes. REST levels shown are 17^th^ percentile (%ile), median, and 83^rd^ %ile.

### Single-nucleotide polymorphisms proximate to the MIR153-1 and MIR153-2 genes

When we queried the dbSNP database, we found a total of 43 validated SNPs within 5kb of the *MIR153-1* or *MIR153-2* genes, out of which at least eight SNPs have functions (Tables 2 and 3). These were compared with selected AD-associated endophenotypes, as described herein.

**Table 2.** Distributions of SNP genotypes by sex and *APOE*ε4 status.

| **rs221301** | **CC** | **CG** | **GG** | **rs6436132** | **GG** | **GT** | **TT** |
| --- | --- | --- | --- | --- | --- | --- | --- |
| Female ε4− | 13 | 109 | 236 | Female ε4− | 208 | 124 | 26 |
| Female ε4+ | 17 | 99 | 201 | Female ε4+ | 175 | 123 | 19 |
| Male ε4− | 20 | 144 | 296 | Male ε4− | 274 | 163 | 23 |
| Male ε4+ | 17 | 126 | 277 | Male ε4+ | 240 | 163 | 17 |
| Assoc gene: *MIR153-2* |  |  |  | Assoc gene: *MIR153-1* |  |  |  |
| Olson's T: 1.264, p = 0.261 |  |  |  | Olson's T: 0.245, p = 0.621 |  |  |  |
| **rs221302** | **AA** | **AG** | **GG** | **rs6459797** | **AA** | **AG** | **GG** |
| Female ε4− | 76 | 183 | 99 | Female ε4− | 67 | 167 | 124 |
| Female ε4+ | 60 | 157 | 100 | Female ε4+ | 50 | 160 | 107 |
| Male ε4− | 87 | 240 | 133 | Male ε4− | 85 | 230 | 145 |
| Male ε4+ | 87 | 216 | 117 | Male ε4+ | 75 | 202 | 143 |
| Assoc gene: *MIR153-2* |  |  |  | Assoc gene: *MIR153-2* |  |  |  |
| Olson's T: 1.523, p = 0.217 |  |  |  | Olson's T: 0.000, p = 0.984 |  |  |  |
| **rs736940** | **AA** | **AG** | **GG** | **rs11887718** | **CC** | **CT** | **TT** |
| Female ε4− | 90 | 176 | 92 | Female ε4− | 12 | 106 | 240 |
| Female ε4+ | 84 | 156 | 77 | Female ε4+ | 12 | 96 | 209 |
| Male ε4− | 115 | 242 | 103 | Male ε4− | 17 | 145 | 298 |
| Male ε4+ | 101 | 206 | 113 | Male ε4+ | 16 | 131 | 273 |
| Assoc gene: *MIR153-2* |  |  |  | Assoc gene: *MIR153-1* |  |  |  |
| Olson's T: 0.009, p = 0.923 |  |  |  | Olson's T: 0.023, p = 0.879 |  |  |  |
| **rs4674374** | **CC** | **CT** | **TT** | **rs17847406** | **CC** | **CG** | **GG** |
| Female ε4− | 13 | 84 | 261 | Female ε4− | 294 | 56 | 8 |
| Female ε4+ | 10 | 84 | 223 | Female ε4+ | 248 | 67 | 2 |
| Male ε4− | 12 | 122 | 326 | Male ε4− | 381 | 76 | 3 |
| Male ε4+ | 13 | 109 | 298 | Male ε4+ | 339 | 79 | 2 |
| Assoc gene: *MIR153-1* |  |  |  | Assoc gene: *MIR153-1* |  |  |  |
| Olson's T: 2.824, p = 0.093 |  |  |  | Olson's T: 0.002, p = 0.961 |  |  |  |

**Table 3.** Endophenotypes affected by *MIR153-1* or *MIR153-2* SNPs.

| Gene | Endophenotype | SNP |
| --- | --- | --- |
| *MIR153-1* | ADAS13 | rs6436132 |
| *MIR153-2* | MMSE | rs6459797 |
| *MIR153-2* | Aβ deposition | rs736940 |
| *MIR153-2* | Entorhinal Cortex Thickness | rs221301 |
| *MIR153-2* | CSF tau | rs221302 |
| *MIR153-2* | RAVLT | rs6459797 |
| *MIR153-1* | CSF Aβ | rs4674374 |
| *MIR153-1* | FDG-PET | rs11887718 |
| *MIR153-1* | Hippocampus Volume | rs17847406 |

### SNPs in the MIR153-1 and MIR153-2 genes influenced levels of eight AD endophenotypes

We tested for associations among the eight SNP genotypes and nine AD endophenotypes, with age, sex, *APOE*ε4 allele presence, and years of formal education as potential covariates of interaction. Competing models were compared via AICc, and the most entropically favorable model for each endophenotype was analyzed by ANOVA and post-hoc comparisons adjusted by BH fdr. Strengths of these associations were assessed via linear models as described herein and expressed as either Nakagawa’s *R*^2^ [99] or as ω^2^ [100]. Models are presented in order from strongest to weakest estimated combined and interactive associations between SNPs and endophenotypes. None of the SNPs were associated with composite scores for memory (data not shown). By contrast, each of the eight SNPs was associated individually with each of the remaining nine AD endophenotypes (Table 3).The majority allele and genotype of rs6436132 near *MIR153-1* is associated with higher ADAS13 scores (data not shown), however, this could be reduced by sex (female lower). Surprisingly, the presence of the *APOE*ε4 allele is associated with lower ADAS13 scores as well. In addition to SNP-associated effects, greater age and lower education are associated with increases in ADAS13.

The minority allele and genotype of rs6459797 is associated with lower MMSE scores (data not shown) in interaction with fewer years of formal education. Although MMSE scores did not differ at highest educational levels, when education was lower, the minority (AA) genotype associated with significant reduction in MMSE. In addition, independently of SNP genotype, increase in age associated with reduced MMSE, as did the presence of at least one *APOE*ε4 allele.

Effects of rs736940 on Aβ deposition are strongly influenced by age and sex (Fig. 2B-G, Table 4). Specifically, increasing age accompanied increased Aβ deposition. This effect is significantly modified (as a difference in the slope of the age effect) by rs736940 genotype, but only for male subjects (Fig. 2E-G). In addition, formal education can further modify this complex effect. Specifically, there appears to be a protective relationship between the “A” allele and Aβ deposition (Table 5). In addition, greater education may enhance such protection. Finally, in all cases, *APOE*ε4 allele presence is associated with greater Aβ deposition.

**Fig. 2.**
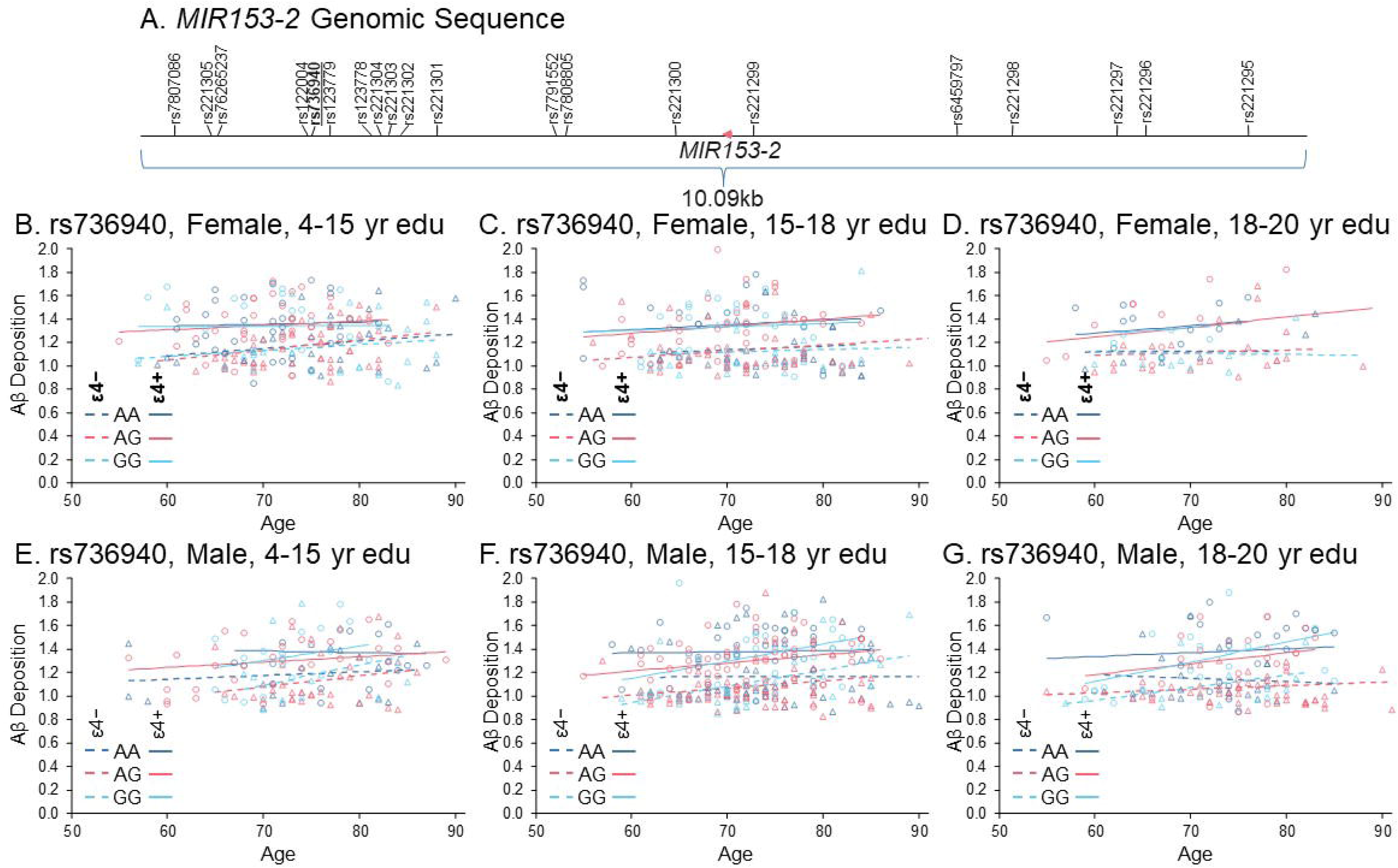
Effects of rs736940 SNP on Aβ deposition. A) 5kb flanking region of the *MIR153-2* gene indicating SNPs. rs736940 in boldface and underlined. Effects are illustrated on the basis of tercile of formal education, in years. B) Effects of SNP and covariates on female subjects with 4-15 years of formal education. C) Effects on female subjects, 15-18 yrs education. D) Effects on female subjects, 18-20 yrs education. E) Effects on male subjects, 4-15 yrs education. F) Effects on male subjects, 15-18 yrs education. G) Effects on male subjects, 18-20 yrs education.

**Table 4.** ANOVA of rs736940 effects on Aβ deposition.

| **Effect** | **F (df,df)** | **p** | **$\omega^2$** |
| --- | --- | --- | --- |
| APOEε4 | 272.44 (1,1033) | < 0.001 | 0.197 |
| Age | 28.95 (1,1033) | < 0.001 | 0.020 |
| rs736940 | 3.77 (2,1033) | 0.023 | 0.004 |
| Education | 1.79 (1,1033) | 0.181 | < 0.001 |
| Sex | 0.96 (1,1033) | 0.328 | < 0.001 |
| Age × rs736940 | 3.28 (2,1033) | 0.038 | 0.003 |
| rs736940 × Sex | 3.10 (2,1033) | 0.045 | 0.003 |
| Age × Sex | 2.77 (1,1033) | 0.096 | 0.001 |
| ε4 × Education | 1.61 (1,1033) | 0.204 | < 0.001 |
| ε4 × Age | 0.41 (1,1033) | 0.522 | < 0.001 |
| Age × Education | 0.12 (1,1033) | 0.728 | < 0.001 |
| Age × rs736940 × Sex | 4.86 (2,1033) | 0.008 | 0.006 |
| ε4 × Age × Educ. | 6.52 (1,1033) | 0.011 | 0.004 |

**Table 5.** Influence of rs736940 genotype on age effect on Aβ deposition.

| Genotype | $\varepsilon_4$ | 4-15 yrs Education | | 15-18 yrs Education | | 18-20 yrs Education | |
| --- | --- | --- | --- | --- | --- | --- | --- |
|  |  | Trend<sup>a</sup> | Group<sup>b</sup> | Trend | Group | Trend | Group |
| AA | – | 0.003 + 0.003 | †‡§ | 0.000 + 0.002 | ‡ | -0.003 + 0.003 | Δ |
| AG | – | 0.009 + 0.003 | †‡ | 0.006 + 0.002 | ‡ | 0.003 + 0.002 | § |
| GG | – | 0.017 + 0.003 | * | 0.014 + 0.003 | *† | 0.011 + 0.003 | †‡ |
| AA | + | -0.001 + 0.003 | § | 0.001 + 0.003 | ‡ | 0.003 + 0.003 | ‡§ |
| AG | + | 0.005 + 0.003 | ‡§ | 0.007 + 0.002 | †‡ | 0.009 + 0.003 | † |
| GG | + | 0.012 + 0.003 | *† | 0.015 + 0.003 | * | 0.017 + 0.003 | * |

<sup>a</sup>Aβ deposition per year of age ± estimated standard error.
<sup>b</sup>Groups determined by pairwise comparisons within age group, adjusted by Benjamini-Hochberg false discovery rate. Conditions sharing symbols do not significantly differ from each other.

We also found significant associations between SNPs in the *MIR153-1* and *MIR153-2* genes and six other endophenotypes, specifically entorhinal cortex thickness, CSF tau levels, RAVLT score, CSF Aβ, brain glucose metabolism, and hippocampus volume (Fig. S1-S6, Tables S1-S3)

### Bioinformatics revealed four potential miR-153-3p sites in the REST 3’-UTR

Multiple bioinformatics searches revealed four putative, adjacent binding sites in the REST (NM005612) 3’-UTR (Table 6, Fig. 3A-E). Each site in the REST 3’-UTR was predicted to have a loop of non-homology between the miR-153-3p and REST sequences. Comparison of the human sequence with the Multiz interspecies alignment showed varying amounts of conservation. Three of these sites had very low conservation (Fig. 3B, D, E) in mice, a commonly used AD research model. The most frequently predicted site (Fig. 3B) shares sequence conservation in the miR-153-3p seed sequence but low conservation in the remainder of the miRNA sequence.

**Table 6.** Predicted sites of miR-153-3p in REST 3’-UTR.

| Utility | Site 1 | Site 2 | Site 3 | Site 4 |
| --- | --- | --- | --- | --- |
| TargetScan<sup>a</sup> | -0.28 | Not predicted | Not predicted | Not predicted |
| DIANA-microT<sup>b</sup> | 0.0178 | 0.0035 | 0.0017 | Not predicted |
| miRDB<sup>c</sup> | 66 | Not predicted | Not predicted | Not predicted |
| miRmap<sup>d</sup> | 93.37 | 83.57 | 72.35 | Not predicted |
| StarMir<sup>e</sup> | 0.667 | 0.593 | 0.417 | 0.350 |

<sup>a</sup>Context++ score<sup>b</sup>miTG score/"Score"<sup>c</sup>Target score<sup>d</sup>miRmap score<sup>e</sup> $\Delta$ G of entire miRNA/mRNA hybrid

**Fig. 3.**
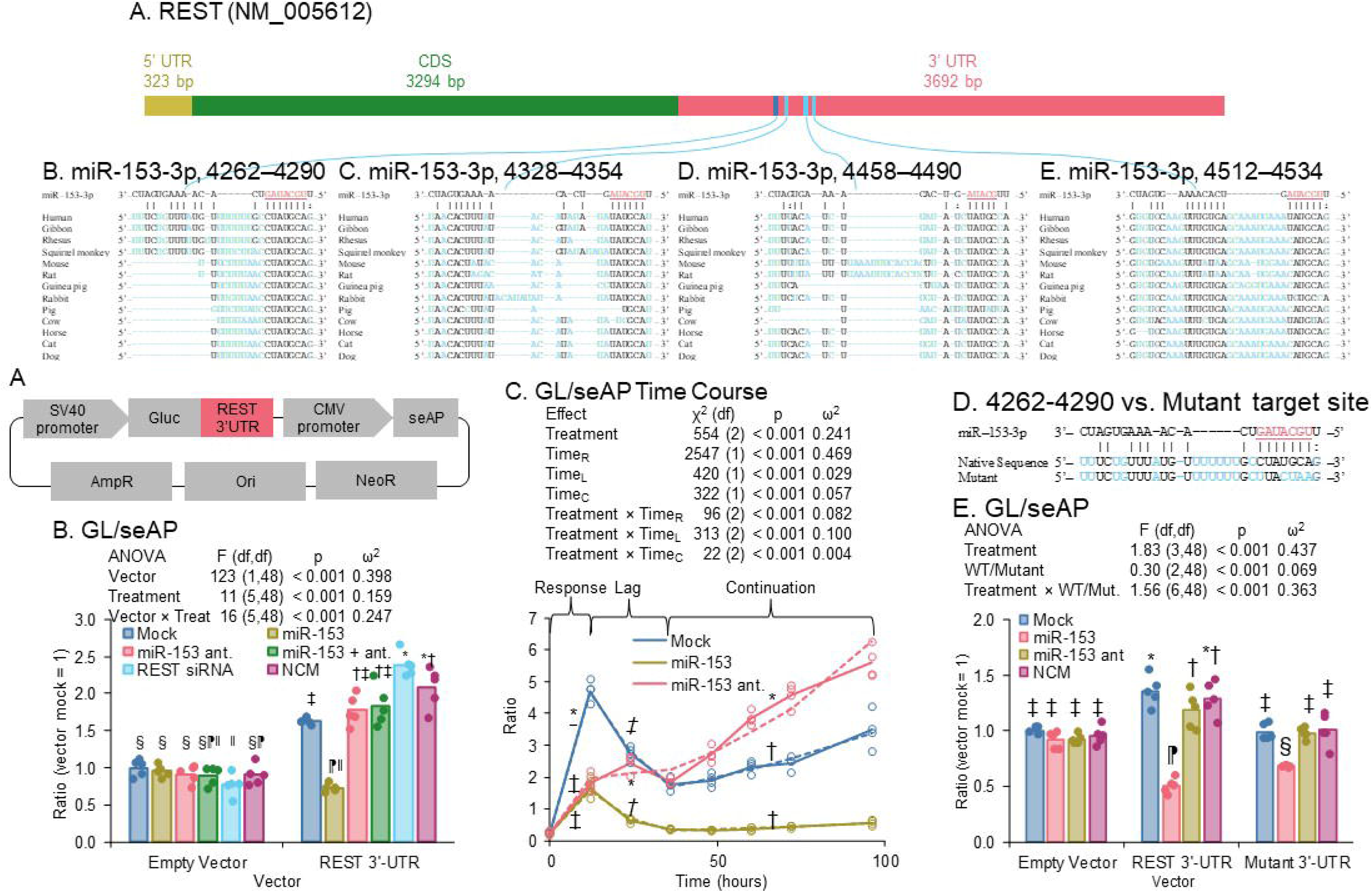
Predicted miR-153-3p sites in REST 3’-UTR and effects of miR-153-3p mimics on REST 3-UTR activity. The REST 3’-UTR (NM005612) was compared with the miR-153-3p sequence via the StarMir utility. Four putative sites were found. Multiz alignments with 11 other mammalian species are shown. A) REST mRNA sequence. B) Site between 4262-4290 (starting from transcription start site). C) Site between 4328-4354. D) Site between 4458-4490. E) Site between 4512-4534. F) Structure of Gluc/seAP vector with REST 3’-UTR. G) Transfection of empty GLuc/seAP vector or vector with REST 3’-UTR, co-transfected with mock, miR-153-3p mimic, miR-153-3p antagomiR, miR-153-3p mimic + antagomiR, REST siRNA, and NCM (negative control), in HeLa cells. miR-153-3p mimic drove reduced expression when REST 3’-UTR was present. H) Time course of effects of miR-153-3p mimic or miR-153-3p antagomir on GLuc/seAP ratio in HeLa cells. Over a period of 36-96 hours, a single transfection of miR-153-3p mimic significantly reduced GLuc/seAP ratio, while a single transfection of the miR-153-3p antagomiR significantly increased such signal. The interval from 36-96 hours was modeled as a repeat-measure ANOVA. I) REST 3’-UTR was mutated as shown. J) Transfection of empty GLuc/seAP vector and vector with native or mutant REST 3’-UTR, co-transfected with mock, miR-153-3p mimic, miR-153-3p antagomiR and NCM (negative control miRNA), in HeLa cells. Symbols indicate differing statistical groups at p ≤ 0.05. Samples sharing symbols did not significantly differ.

### miR-153-3p targeted and reduced REST 3’-UTR activities

We cloned the REST 3’-UTR into the GLuc/seAP vector (Fig. 3F), transfected into HeLa cells, and treated with miR-153-3p mimics and antagomiRs, as well as REST siRNA. We found that miR-153-3p significantly reduced GL/seAP signal (Fig. 3G), which was reversed by co-treatment with an miR-153-3p antagomiR. As expected, the commercial REST siRNA did not produce a significant difference from mock treatment, since it targets the REST mRNA protein coding sequence (CDS). We also noticed that REST 3’-UTR containing vector transfection alone had a higher GL/SEAP ratio compared to empty vector.

We also transfected HeLa cells with the REST/GLuc fused plasmid and treated in parallel with miR-153-3p or an miR-153-3p antagomiR and measured secreted GL/seAP activities at intervals from transfection to 96 hours (Fig. 3H). We found that slopes of each treatment could significantly differ according to phases we designated as “response” (R), “lag” (L)”, or “continuation” (C). Specifically, in the R phase, mock treatment significantly differed from both miR-153-3p mimic and antagomiR. In the L phase, each slope significantly differed from the others. In the C phase, levels of signal significantly continued to increase vs. both mock treatment and miR-153-3p mimic treatment. Note that this analysis refers to the *slopes* of the responses, not levels at individual time points. Activity of exogenous miR-153-3p lasted at least 96 hours after treatment, while antagomiR significantly blocked endogenous miR-53-3p for the same time frame.

### Mutational analyses confirm one miR-153-3p functional target site in the REST 3’-UTR

Four sites were predicted to bind on REST 3’-UTR, but only one site (Table 6) was predicted by all bioinformatic tools we used. Based on the prediction, a mutated plasmid on the seed sequence-binding region within the highest scored site was generated and applied to confirm functional interaction between miR-153-3p and REST 3’-UTR. Five out of seven nucleotides in the seed sequence binding region were mutated (Fig. 3I). The mutant partially reversed the inhibitory effects of miR-153-3p. However, activity of the mutant REST 3’-UTR clone was still significantly reduced by miR-153-3p (Fig. 3J), implying that miR-153-3p may also function through one or more of the other predicted sites on the REST 3’-UTR. In addition, abolition of this site significantly reduced GL/seAP signal in mock-transfected mutant vs. wildtype 3’-UTR clones. Thus, although more than one site may interact with miR-153-3p, this particular site may function to maintain higher levels of REST through some other mechanism.

### Bioinformatic search revealed two potential miR-153-3p sites in the SNCA 3’-UTR

A search with the StarMir utility revealed two putative miR-153-3p binding sites in the SNCA (NM001375287) 3’-UTR (Fig. 4A-C). Comparison of the human sequence with the Multiz interspecies alignment showed consistent conservation *except for mice*. The downstream site was essentially absent in the mouse sequence, and both mouse and rat sequences had a 5-base gap when aligned to the human sequence.

**Fig. 4.**
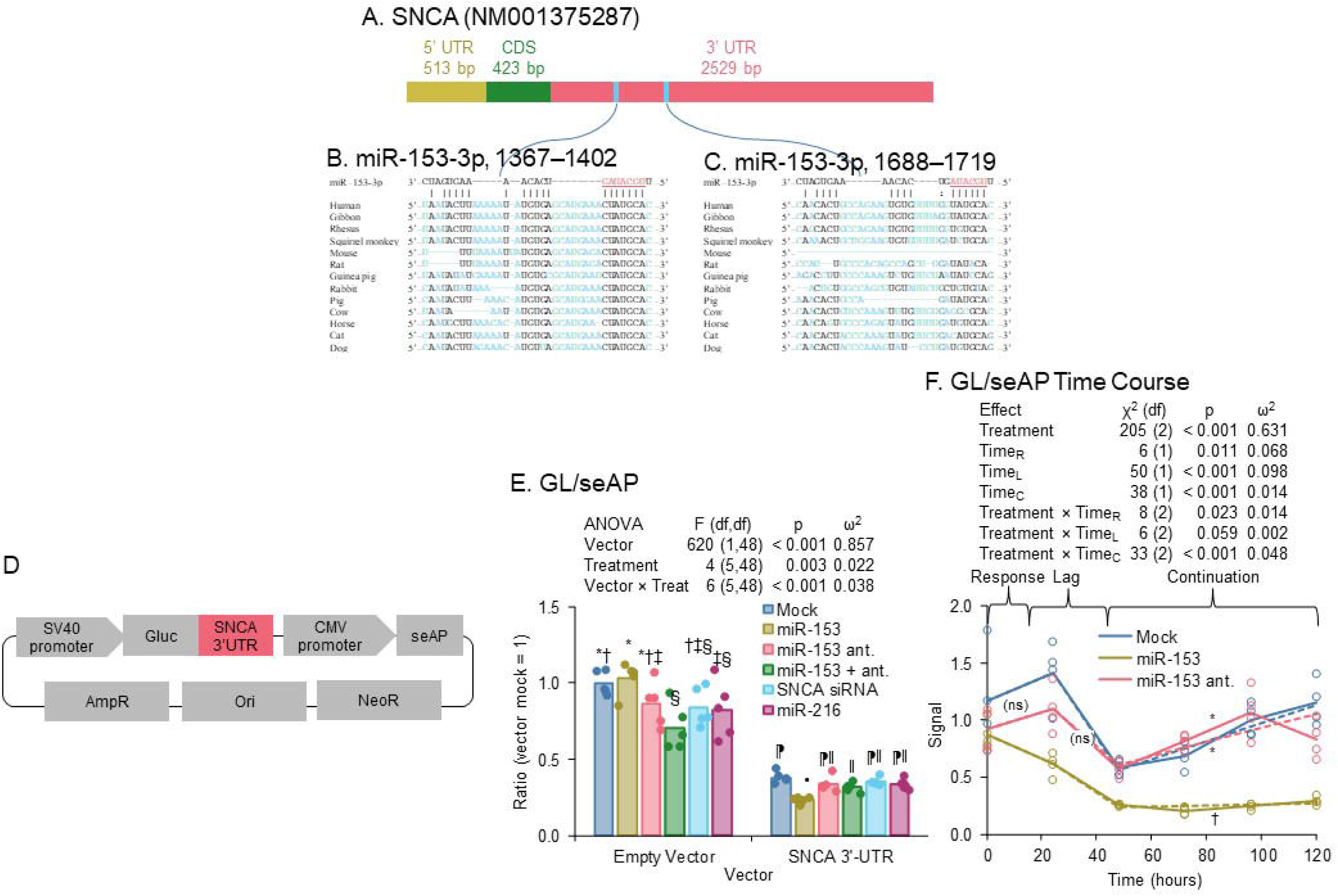
Predicted miR-153-3p sites in SNCA 3’-UTR. GLuc/seAP assay of transfected clones vs. miR-153-3p and associated oligomers. The SNCA 3’-UTR (NM001375287) was compared with the miR-153-3p sequence via the StarMir utility. Two putative sites were found. Multiz alignments with 11 other mammalian species are shown. A) SNCA mRNA sequence. B) Site between 1367-1402 (starting from transcription start site). C) Site between 1688-1719. D) GLuc/seAP vector with SNCA 3’-UTR insert. E) Transfection of empty GLuc/seAP vector or vector with SNCA 3’-UTR, co-transfected with mock, miR-153-3p mimic, miR-153-3p antagomiR, miR-153-3p mimic + antagomiR, SNCA siRNA, and miR-216 (alleged negative control), in HeLa cells. miR-153-3p mimic drove reduced expression when SNCA 3’-UTR was present. F) Time course of effects of miR-153-3p mimic or miR-153-3p antagomir on GLuc/seAP ratio in HeLa cells. Over a period of 48-120 hours, a single transfection of miR-153-3p mimic significantly reduced GLuc/seAP ratio. Symbols indicate differing statistical groups at p ≤ 0.05. Samples sharing symbols did not significantly differ.

### miR-153-3p targeted and reduced SNCA 3’-UTR activities

When we cloned the SNCA 3’-UTR into the GLuc/seAP vector, transfected into HeLa cells, and treated with miR-153-3p mimics and antagomiRs and SNCA siRNA, we found that miR-153-3p significantly reduced GL/seAP signal when transfected along with the SNCA fusion vector (Fig. 4E), which was reversed by co-treatment with an miR-153-3p antagomiR. As expected, the commercial SNCA siRNA did not produce a significant difference from mock treatment, since it targeted the SNCA mRNA CDS.

We also transfected HeLa cultures with the SNCA/GLuc fusion and treated in parallel with miR-153-3p or the miR-153-3p antagomiR and measured secreted GL/seAP activities at intervals from transfection to 120 hours (Fig. 4F). We found that the slope of the miR-153-3p treatment significantly differed from each other slope during the C phase. No significant differences in slopes appeared in the R or L phases.

### Bioinformatics revealed one potential miR-153-3p site in the APP 3’-UTR

A search with the StarMir utility revealed one putative binding site in the APP (NM000484) 3’-UTR (Fig. 5A). Comparison of the human sequence with the Multiz interspecies alignment showed high conservation (Fig. 5B).

**Fig. 5.**
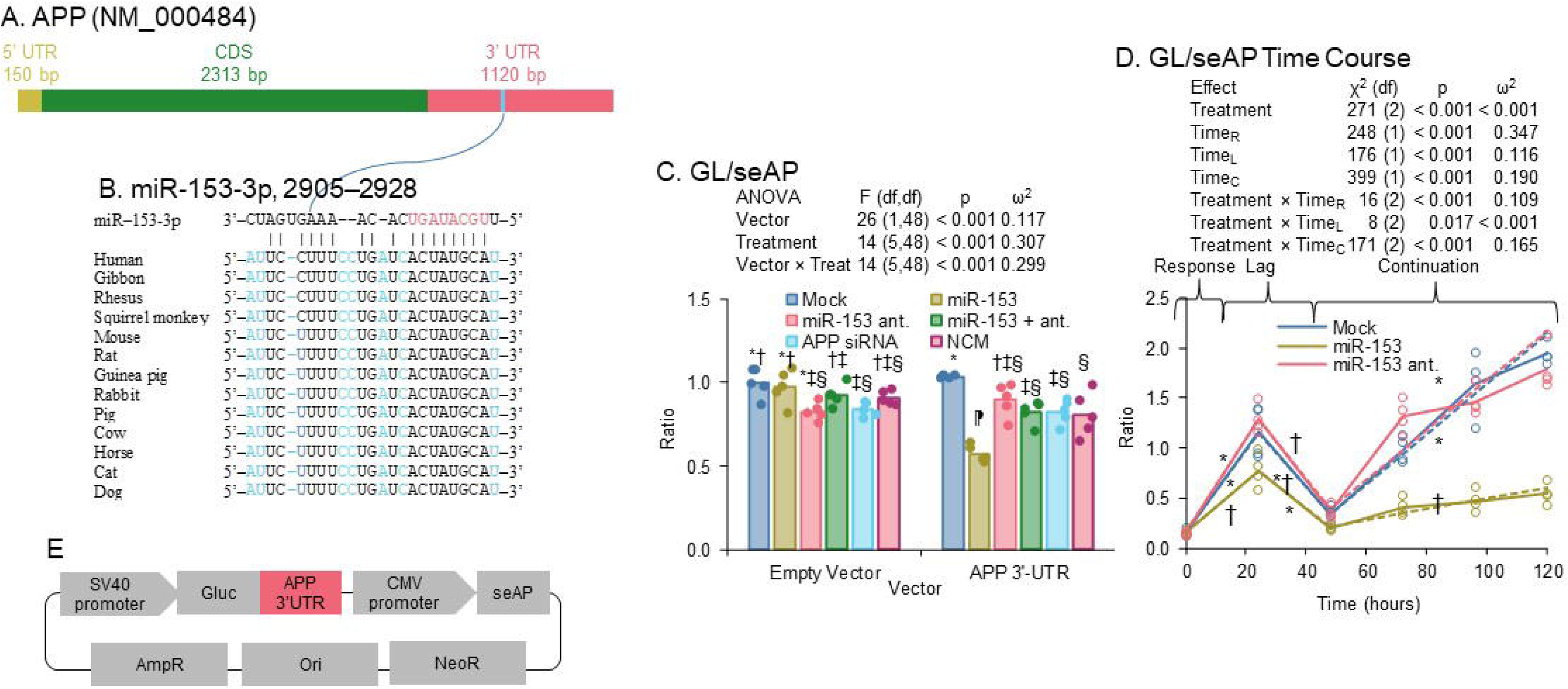
Predicted miR-153-3p sites in APP 3’-UTR. GLuc/seAP assay of transfected clones vs. miR-153-3p and associated oligomers. The APP 3’-UTR (NM000484) was compared with the miR-153-3p sequence via the StarMir utility. One putative site was found. Multiz alignments with 11 other mammalian species are shown. A) APP mRNA sequence. B) Site between 2905-2928 (starting from transcription start site). C) Site between 2905-2928. D) Transfection of empty GLuc/seAP vector or vector with APP 3’-UTR, co-transfected with mock, miR-153-3p mimic, miR-153-3p antagomiR, miR-153-3p mimic + antagomiR, APP siRNA, and NCM (negative control), in HeLa cells. miR-153-3p mimic drove reduced expression when APP 3’-UTR was present. E) Time course of effects of miR-153-3p mimic or miR-153-3p antagomir on GLuc/seAP ratio in HeLa cells. Over a period of 48-120 hours, a single transfection of miR-153-3p mimic significantly reduced GLuc/seAP ratio Symbols indicate differing statistical groups at p ≤ 0.05. Samples sharing symbols did not significantly differ. E) Diagram of GLuc/seAP vector with APP 3’-UTR.

### miR-153-3p targeted and reduced APP 3’-UTR activities

When we cloned the APP 3’-UTR into the GLuc/seAP vector, transfected into HeLa cells, and treated with miR-153-3p mimics and antagomiRs, and APP siRNA, we found that miR-153-3p significantly reduced GL/seAP signal when transfected along with the APP fusion vector (Fig. 5C), which was reversed by co-treatment with an miR-153-3p antagomiR. As expected, APP siRNA did not produce a significant difference from Mock treatment, since it targets the APP mRNA CDS.

We also transfected HeLa cultures with the APP/GLuc fusion and treated in parallel with miR-153-3p or the miR-153-3p antagomiR and measured secreted GL/seAP activities at intervals from transfection to 120 hours (Fig. 5D). We found that the slopes of the miR-153-3p treatment significantly differed from each other slope in the R and C phases. Treatment with miR-153-3p mimic had a less steep slope than both mock and miR-153-3p antagomiR.

### miR-153-3p affected REST, APP and SNCA expression levels

We treated HeLa and Diff-NB cells with miR-153-3p, miR-153-3p antagomiR, miR-153-3p + antagomiR, SNCA siRNA, and miR216 which does not have any prediction on our target proteins of interest. (Fig. 6) Notably, miR-153-3p treatment significantly reduced REST and APP protein levels and its effects were reversed by co-treatment with miR-153-3p antagomiR in both cell lines. SNCA levels were reduced only in differentiated neuronal cells. Surprisingly, treatment by SNCA siRNA significantly elevated both REST and APP levels in neuronal cells. In addition, miR216 elevated SCNA levels, perhaps through an indirect or non-canonical mechanism

**Fig. 6.**
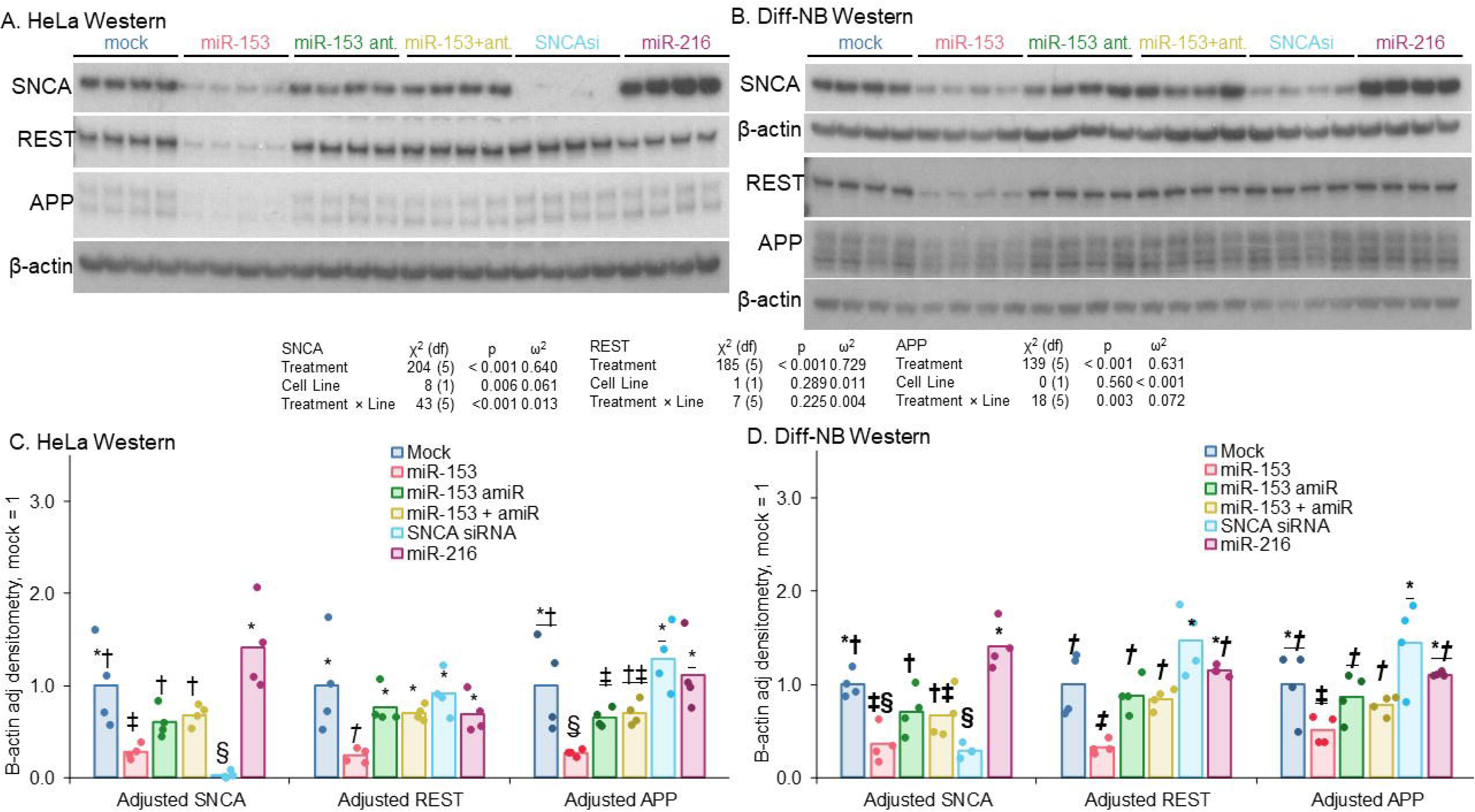
Effects of miR-153-3p treatment in HeLa and differentiated neuronal cells. Cultures of A) HeLa and B) NB were treated with miR-153-3p mimic, antagomiR of miR-153-3p, miR-153-3p mimic plus antagomiR, siRNA vs. SNCA, and miR-216 (alleged negative control), and levels of SNCA, REST, and APP measured. Densitometry revealed that in C) HeLa and D) Diff-NB, miR-153-3p mimic consistently reduced SNCA, REST, and APP protein signal (adjusted by β-actin signal). Of potential note, treatment by miR-216 increased levels of SNCA vs. mock treatment. Symbols indicate differing statistical groups at p ≤ 0.05. Samples sharing symbols did not significantly differ.

### miR-153-3p altered neuronal differentiation

To study potential effects of miR-153-3p on neuronal differentiation via REST regulation, we transfected neuronal stem cells with combinations of miR-153-3p, antagomiRs, and REST siRNA. Treatment by miR-153-3p reduced REST expression in neuronal stem cells, while also significantly reducing nestin and increasing doublecortin protein levels. (Fig. 7A, C) REST knockdown by siRNA also reduced nestin protein levels but did not significantly alter doublecortin levels.

**Fig. 7.**
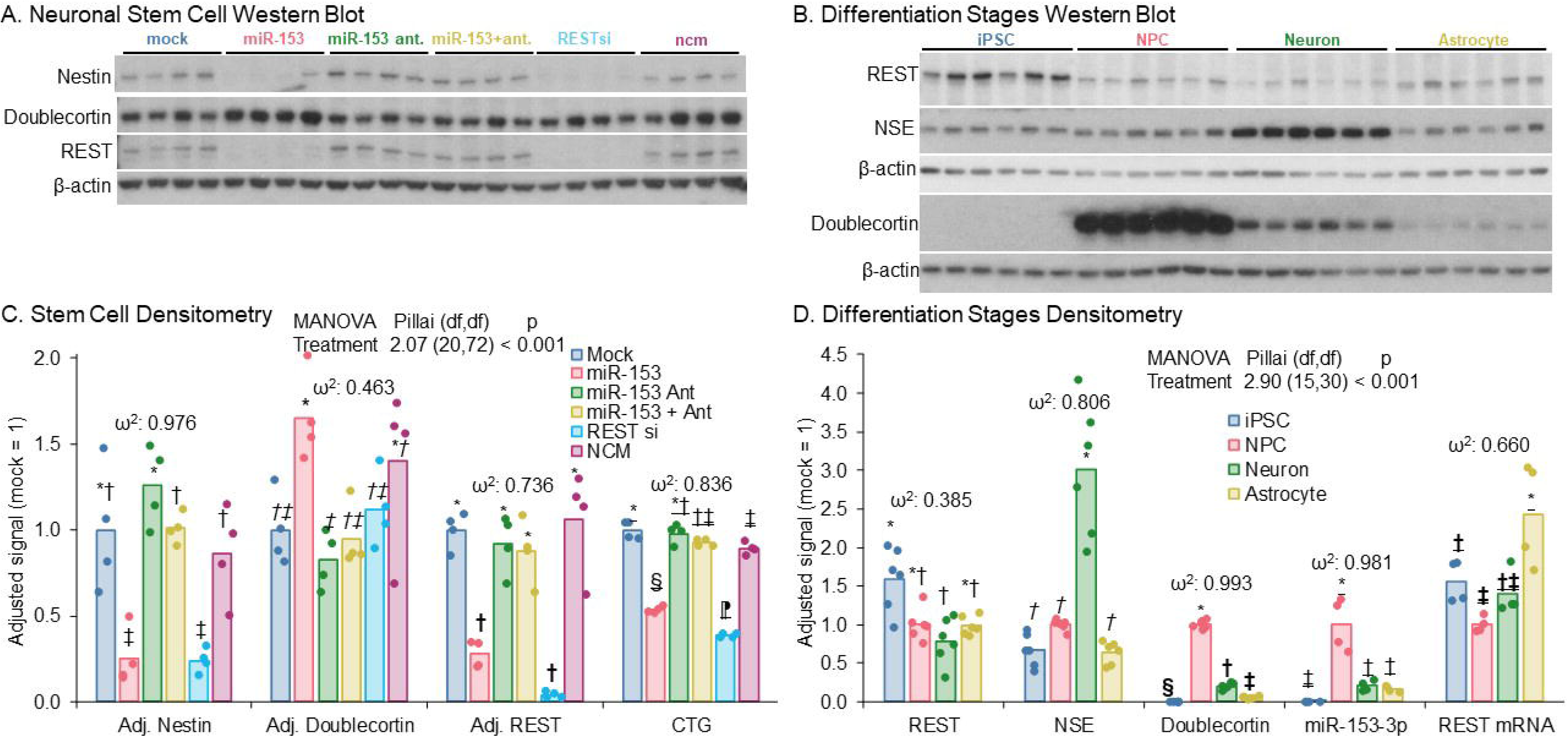
Effects of miR-153-3p mimic treatments on REST, nestin and doublecortin in neural stem cells and variation of effect by differentiation. Neural stem cells were cultured and treated with miR-153-3p mimic, and other oligomers as described in the main text. In addition, human iPS cells were differentiated into neural progenitor cells (npc), neurons and astrocytes. A) Western blot. B) Western blot of REST, NSE, doublecortin and β-actin proteins. C) Densitometry of REST, nestin and doublecortin proteins, as well as levels of CTG. Treatment with miR-153 and REST siRNA significantly associated with reductions in REST, nestin protein and CTG. Treatment with miR-153 but not REST siRNA increased doublecortin protein. D) REST mRNA levels of four different stages of cells; Densitometric quantification of REST, NSE and doublecortin proteins; endogenous miR-153 levels of ipsc, npc, neurons and astrocytes. Symbols indicate differing statistical groups at p ≤ 0.05. Samples sharing symbols did not significantly differ.

### Neuronal differentiation alters levels of miR-153-3p and REST, as well as NSE and doublecortin

Human iPSC cells were differentiated into neural progenitor cells (NPC), neurons and astrocytes. Levels of miR-153-3p, REST mRNA, REST protein, NSE, and doublecortin were measured (Fig. 7B, D). miR-153-3p levels were highest in NPC cells and lowest in naïve iPSC cells. In addition, both neurons and astrocytes have elevated miR-153-3p vs. iPSC. REST mRNA and protein have different patterns of expression from each other; specifically, REST protein is highest in iPSC and lowest in neurons, although the signals seen in NPCs and astrocytes are not significantly higher than in neurons. On the other hand, the highest levels of REST mRNA were seen in astrocytes, and all other differentiation stages were in the same lower group by statistical significance. Doublecortin protein levels fairly closely matched those of miR-153-3p. Finally, levels of NSE were apparently unrelated to other markers measured, with the highest levels seen in neurons.

### miR-153-3p is important for axonal guidance

To determine the overall impact of miR-153-3p, differentiated neuroblastoma cells were transfected with miR-153-3p, its antagomiR or mock and subjected to RNA-seq analysis. KEGG pathway enrichment assay (Fig. 8) showed that gene expression altered by miR-153-3p is enriched in axonal guidance, cell cycle, and focal adhesion.

**Fig. 8.**
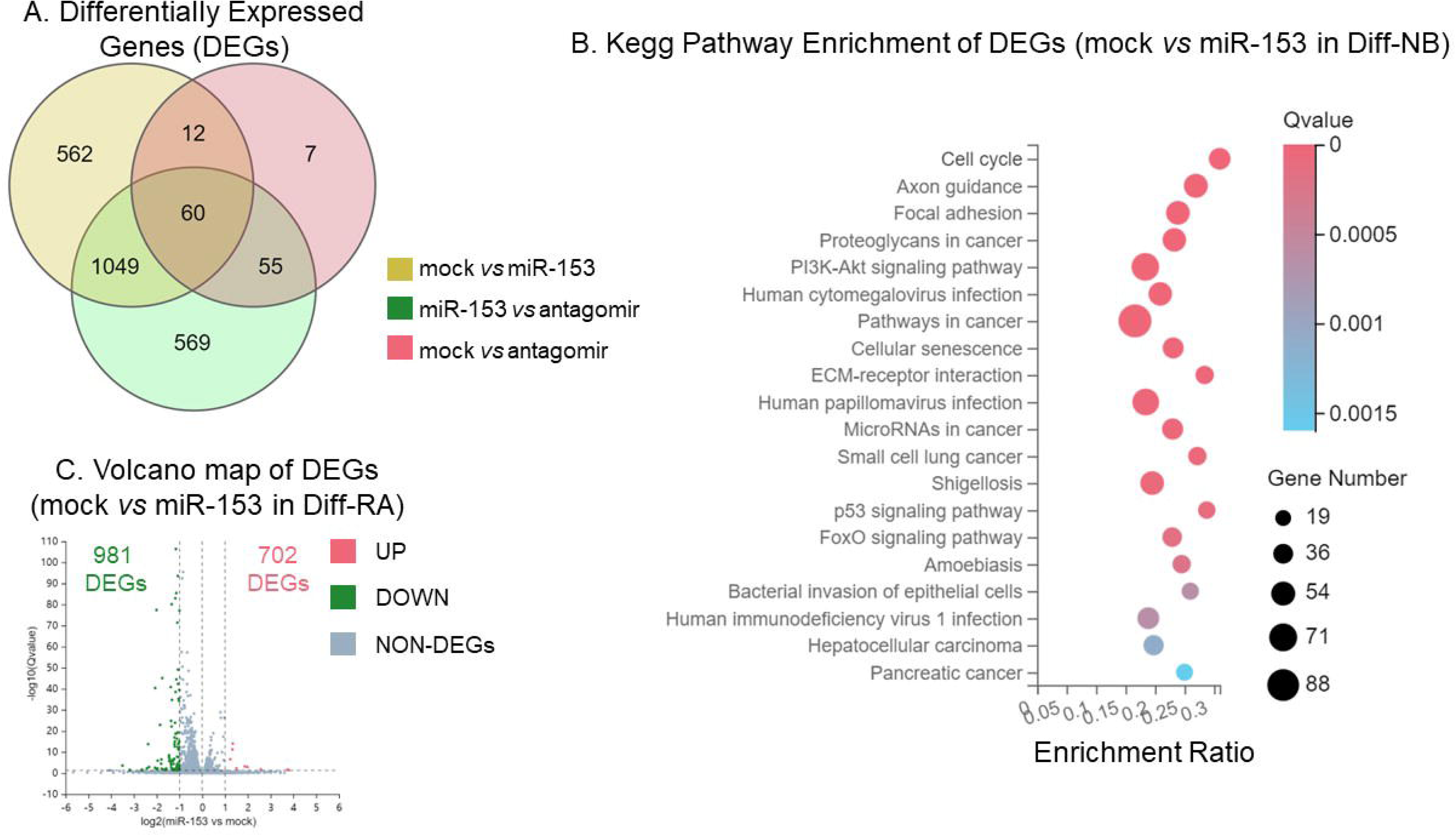
RNA-sequencing data analysis of miR-153 transfection in Diff-NB cells. Diff-NB cells were transfected with miR-153 or miR-153 antigomiR. mRNA samples were subjected to RNA-seq assay followed by pathway analysis. A) Count of differentially expressed genes. B) Kegg pathway enrichment, mock vs. miR-153-3p mimic. C) Volcano map of differentially expressed genes.

### miR-153-3p-specific proteome alterations in mixed human brain cultures

MT proteomic analysis identified 316 significantly differentially expressed proteins in miR-153-3p compared to mock-transfected Diff-NB cultures, whereas 167 proteins were altered by miR-153-3p antagomir treatments (data not shown). Overall, 115 proteins from the combined miR-153-3p and the antagomiR datasets were inversely related to each other with a base-2 log effect difference (miR-153-3p÷mock − antagoMir÷mock) of at least ±0.2, and a p value ≤ 0.05 (Fig. 9).

**Fig. 9.**
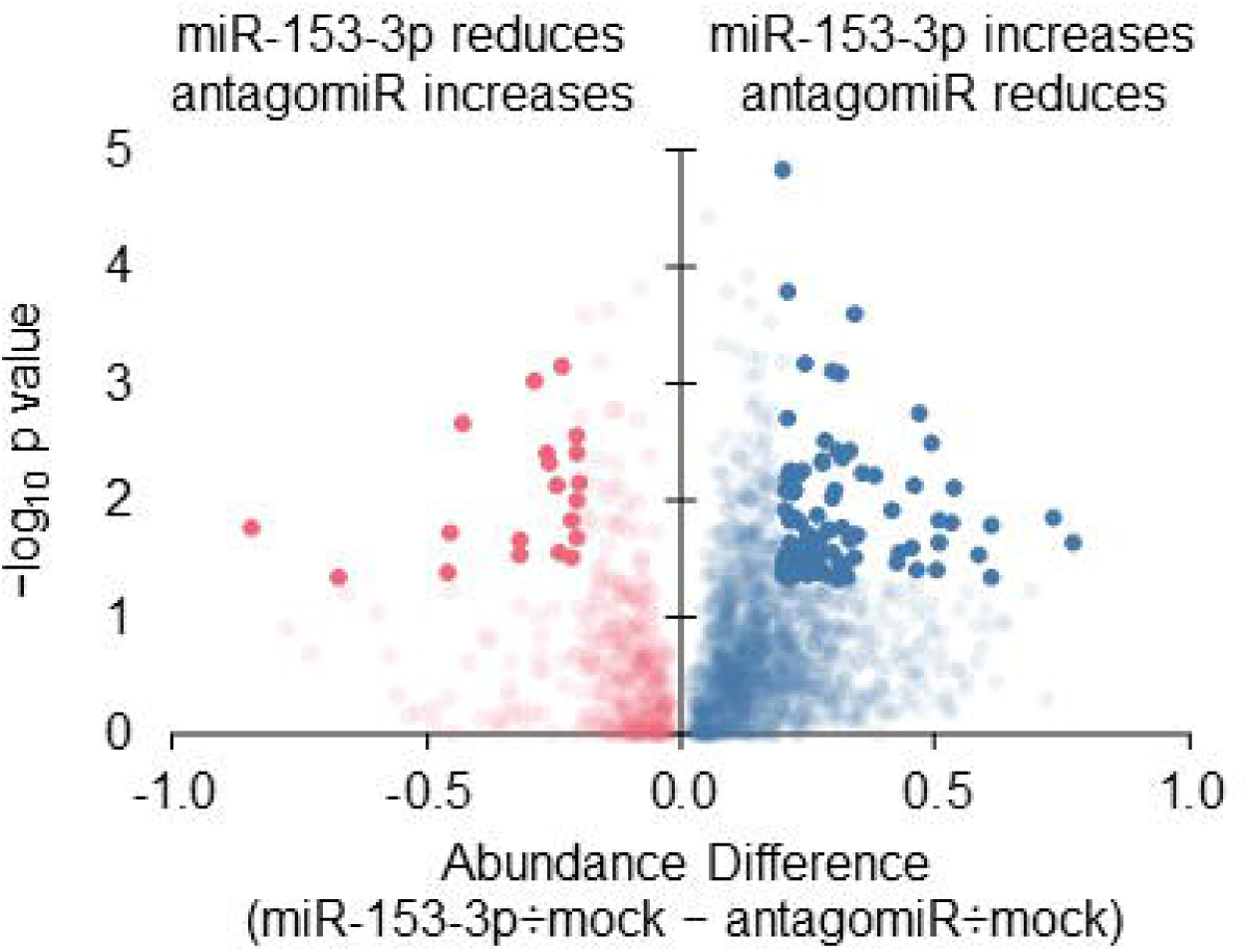
Proteomic abundance outcomes. Proteomic analysis was performed, and data filtered as described herein. Red points represent proteins with miR-153-3p mimic ÷ mock abundance < 1 and antagomR ÷ mock abundance > 1. Blue points represent proteins with miR-153-3p mimic ÷ mock abundance > 1 and antagomR ÷ mock abundance < 1. Solid points have log_2_ abundance absolute differences of ≥ 0.2 and p ≤ 0.05.

### miR-153-3p-related interactomes in hippocampus and frontal cortex

We generated protein-protein interaction networks from proteomic analysis of cells treated with miR-153-3p and its corresponding antagomir, along with APP, DCX, NES, REST, and SNCA, for which we directly measured miR-153-3p activity in cell cultures. We also created protein-protein interaction networks for human hippocampus and frontal cortex (data not shown). Interaction strengths were estimated via STRING, along with reporting enriched gene ontology, reactome, and human disease phenotype pathways. Those pathways with at least 15-fold strength over what would be expected for a random network are shown (Tables 7-8). Full reports can be found at https://string-db.org/cgi/network?taskId=bQXeAn38gHN5 for hippocampus and https://string-db.org/cgi/network?taskId=bocxSsRn3X5b for frontal cortex.

**Table 7.** Selected enriched hippocampus network associations.

| Enrichment<sup>a</sup> | Description | Count<sup>b</sup> |  | Str<sup>c</sup> | fdr<sup>d</sup> | Network Proteins<sup>e</sup> |
| --- | --- | --- | --- | --- | --- | --- |
|  |  | obs | bck |  |  |  |
| Component | NELF complex | 2 | 4 | 58 | 0.030 | RDBP, WHSC2 |
| Component | Beta-catenin destruction complex | 3 | 11 | 31 | 0.008 | APC, SIAH1, CTNNB1 |
| Component | RNA polymerase II transcription repressor complex | 3 | 15 | 23 | 0.015 | DDX20, MYC, CTNNB1 |
| Component | Apicolateral plasma membrane | 3 | 22 | 15 | 0.033 | <u>CXADR</u>, NEDD4, CTNNB1 |
| Phenotypes | Pilomatrixoma | 3 | 6 | 58 | 0.046 | APC, CREBBP, CTNNB1 |
| Phenotypes | Micrographia | 3 | 7 | 49 | 0.046 | <u>VPS13A</u>, PARK2, **SNCA** |
| Phenotypes | Scissor gait | 3 | 9 | 38 | 0.048 | SHMT2, <u>ATP6AP2</u>, **SNCA** |
| Phenotypes | Low hanging columella | 4 | 26 | 18 | 0.047 | CREBBP, <u>KDM3B</u>, CTNNB1, PQBP1 |
| Process | miRNA-mediated gene silencing by mRNA destabilization | 2 | 3 | 76 | 0.045 | EIF2C3, MOV10 |
| Process | Positive regulation of intrinsic apoptotic signaling pathway by p53 class mediator | 3 | 7 | 49 | 0.008 | <u>EIF5A</u>, UBB, MYC |
| Process | Negative regulation of monooxygenase activity | 3 | 12 | 29 | 0.022 | VDR, <u>ENG</u>, **SNCA** |
| Process | Spliceosomal tri-snRNP complex assembly | 3 | 13 | 26 | 0.026 | DDX20, <u>LSM2</u>, <u>SRSF12</u> |
| Process | Positive regulation of pseudopodium assembly | 3 | 13 | 26 | 0.026 | APC, CDC42EP2, CDC42 |
| Process | Layer formation in cerebral cortex | 3 | 15 | 23 | 0.032 | CDK5, **DCX**, CTNNB1 |
| Process | Synaptic transmission, dopaminergic | 3 | 16 | 21 | 0.037 | PARK2, CDK5, **SNCA** |
| Process | Lipoprotein transport | 3 | 17 | 20 | 0.041 | <u>UNC119</u>, <u>UNC119B</u>, ZDHHC17 |
| Process | Receptor catabolic process | 3 | 18 | 19 | 0.046 | <u>PIK3R4</u>, CDK5, NEDD4 |
| Process | Synaptic vesicle transport | 5 | 33 | 17 | 0.003 | <u>AP3M1</u>, CDK5, LIN7A, CTNNB1, **SNCA** |
| Process | Cerebral cortex radial glia-guided migration | 4 | 27 | 17 | 0.014 | DISC1, CDK5, **DCX**, CTNNB1 |
| Process | Protein targeting to lysosome | 4 | 28 | 16 | 0.015 | HGS, <u>AP3M1</u>, <u>PIK3R4</u>, NEDD4 |
| Process | Negative regulation of oxidoreductase activity | 4 | 28 | 16 | 0.015 | VDR, PARK2, <u>ENG</u>, **SNCA** |
| Reactome | LRR FLII-interacting protein 1 (LRRFIP1) activates type I IFN production | 3 | 5 | 69 | 0.004 | CREBBP, <u>LRRFIP1</u>, CTNNB1 |
| Reactome | Translation of Replicase and Assembly of the Replication Transcription Complex | 3 | 13 | 26 | 0.018 | <u>PIK3R4</u>, <u>CHMP4A</u>, MAP1LC3B |
| Reactome | Regulation of NPAS4 gene expression | 3 | 13 | 26 | 0.018 | **REST**, EIF2C3, MOV10 |
| Reactome | Translation of Replicase and Assembly of the Replication Transcription Complex | 3 | 14 | 25 | 0.019 | <u>PIK3R4</u>, <u>CHMP4A</u>, MAP1LC3B |
| Reactome | Listeria monocytogenes entry into host cells | 4 | 20 | 23 | 0.005 | HGS, SH3GL2, UBB, CTNNB1 |
| Reactome | InlB-mediated entry of Listeria monocytogenes into host cell | 3 | 15 | 23 | 0.020 | HGS, SH3GL2, UBB |
| Reactome | PINK1-PRKN Mediated Mitophagy | 4 | 22 | 21 | 0.006 | MTERFD1, PARK2, UBB, MAP1LC3B |
| Reactome | EGFR downregulation | 5 | 31 | 19 | 0.002 | EGFR, HGS, SH3GL2, UBB, CDC42 |
| Reactome | Transcriptional Regulation by NPAS4 | 5 | 34 | 17 | 0.003 | CREBBP, **REST**, EIF2C3, MOV10, CDK5 |
| Reactome | Negative regulation of MET activity | 3 | 21 | 16 | 0.038 | HGS, SH3GL2, UBB |

<sup>a</sup>Enrichment types: Component: Cellular component; Phenotype: Human phenotype; Process: Biological process; Reactome: Reactome pathway.
<sup>b</sup>Gene Count; obs: Observed; bck: Background.
<sup>c</sup>Str: Fold-strength, observed ÷ expected. Str 10 = ten times greater incidence than expected.
<sup>d</sup>fdr: Benjamini-Hochberg false discovery rate
<sup>e</sup>**Boldface**: mir-153-3p activity directly confirmed in cell culture. Italic underline: Responder to miR-153-3p by proteomics.
Unmodified: Produced by NetworkAnalyst.

**Table 8.** Selected enriched frontal cortex network associations.

| Enrichment<sup>a</sup> | Description | Count<sup>b</sup> |  | Str<sup>c</sup> | fdr<sup>d</sup> | Network Proteins<sup>e</sup> |
| --- | --- | --- | --- | --- | --- | --- |
|  |  | obs | bck |  |  |  |
| Component | NELF complex | 3 | 4 | 83 | 0.002 | COBRA1, RDBP, WHSC2 |
| Component | U2-type catalytic step 2 spliceosome | 5 | 30 | 19 | 0.001 | <u>AQR</u>, PRPF19, <u>CDC40</u>, <u>CWC22</u>, SNRPB |
| Function | Phosphoserine residue binding | 3 | 9 | 37 | 0.047 | PIN1, YWHAB, NEDD4 |
| Phenotype | Micrographia | 3 | 7 | 48 | 0.025 | <u>VPS13A</u>, PARK2, **SNCA** |
| Phenotype | Scissor gait | 3 | 9 | 37 | 0.038 | SHMT2, <u>ATP6AP2</u>, **SNCA** |
| Phenotype | Abnormality of the temporomandibular joint | 3 | 11 | 30 | 0.050 | <u>COL1A1</u>, <u>VPS13A</u>, <u>COL5A1</u> |
| Process | miRNA-mediated gene silencing by mRNA destabilization | 2 | 3 | 74 | 0.043 | EIF2C3, MOV10 |
| Process | Negative regulation of protein export from nucleus | 3 | 6 | 56 | 0.006 | BARD1, <u>TXN</u>, CDK5 |
| Process | Aggresome assembly | 3 | 6 | 56 | 0.006 | PARK2, BAG3, HDAC6 |
| Process | Collateral sprouting | 3 | 12 | 28 | 0.021 | **APP**, HDAC6, BDNF |
| Process | Spliceosomal tri-snRNP complex assembly | 3 | 13 | 26 | 0.025 | PRPF19, <u>LSM2</u>, <u>SRSF12</u> |
| Process | Positive regulation of pseudopodium assembly | 3 | 13 | 26 | 0.025 | APC, CDC42EP2, CDC42 |
| Process | Cellular response to interleukin-17 | 3 | 13 | 26 | 0.025 | TRAF2, <u>STAT3</u>, TRAF6 |
| Process | Synaptic transmission, dopaminergic | 3 | 16 | 21 | 0.036 | PARK2, CDK5, **SNCA** |
| Process | Lipoprotein transport | 3 | 17 | 20 | 0.040 | <u>UNC119</u>, <u>UNC119B</u>, ZDHHC17 |
| Process | Receptor catabolic process | 3 | 18 | 19 | 0.045 | <u>PIK3R4</u>, CDK5, NEDD4 |
| Process | Negative regulation of transcription elongation from RNA polymerase II promoter | 3 | 18 | 19 | 0.045 | COBRA1, RDBP, WHSC2 |
| Process | Negative regulation of oxidoreductase activity | 4 | 28 | 16 | 0.015 | PARK2, <u>ENG</u>, HDAC6, **SNCA** |
| Reactome | LRR FLII-interacting protein 1 (LRRFIP1) activates type I IFN production | 2 | 5 | 45 | 0.027 | CREBBP, <u>LRRFIP1</u> |
| Reactome | Activated NTRK2 signals through CDK5 | 2 | 6 | 37 | 0.033 | BDNF, CDK5 |
| Reactome | Neurofascin interactions | 2 | 7 | 32 | 0.040 | <u>CNTNAP1</u>, **DCX** |
| Reactome | TRAF6 mediated IRF7 activation in TLR7/8 or 9 signaling | 3 | 12 | 28 | 0.009 | TRAF6, UBC, MYD88 |
| Reactome | Competing endogenous RNAs (ceRNAs) regulate PTEN translation | 2 | 8 | 28 | 0.045 | EIF2C3, MOV10 |
| Reactome | p75NTR recruits signalling complexes | 3 | 13 | 26 | 0.009 | TRAF6, UBC, MYD88 |
| Reactome | Regulation of NPAS4 gene expression | 3 | 13 | 26 | 0.009 | **REST**, EIF2C3, MOV10 |
| Reactome | Transcriptional Regulation by NPAS4 | 6 | 34 | 20 | 0.000 | CREBBP, **REST**, EIF2C3, MOV10, BDNF, CDK5 |
| Reactome | Regulation of NF-kappa B signaling | 3 | 17 | 20 | 0.016 | TRAF2, TRAF6, UBC |
| Reactome | Abortive elongation of HIV-1 transcript in the absence of Tat | 4 | 23 | 19 | 0.004 | <u>POLR2J</u>, COBRA1, RDBP, WHSC2 |
| Reactome | TRAF6 mediated NF-kB activation | 4 | 25 | 18 | 0.005 | TRAF2, **APP**, HMGB1, TRAF6 |
| Reactome | PINK1-PRKN Mediated Mitophagy | 3 | 22 | 15 | 0.024 | MTERFD1, PARK2, UBC |
| Reactome | NPAS4 regulates expression of target genes | 3 | 22 | 15 | 0.024 | CREBBP, BDNF, CDK5 |

<sup>a</sup>Enrichment types: Component: Cellular component; Phenotype: Human phenotype; Process: Biological process; Reactome: Reactome pathway.
<sup>b</sup>Gene Count; obs: Observed; bck: Background.
<sup>c</sup>Str: Fold-strength, observed ÷ expected. Str 10 = ten times greater incidence than expected.
<sup>d</sup>fdr: Benjamini-Hochberg false discovery rate
<sup>e</sup>**Boldface:** mir-153-3p activity directly confirmed in cell culture. Italic underline: Responder to miR-153-3p by proteomics.
Unmodified: Produced by NetworkAnalyst.

### Levels of native miR-153-3p differ by cell type

We isolated RNA from epithelial (HeLa), astrocytic (U373) and neuroblastoma (SH-SY5Y and SK-N-SH) cells and from neuron typic SK-N-SH cells treated with retinoic acid and measured levels of miR-153-3p by qRT-PCR. The neuroblastoma-derived cell types had significantly higher basal levels of miR-153-3p than cells of non-neuronal origin (Fig. 10A).

**Fig. 10.**
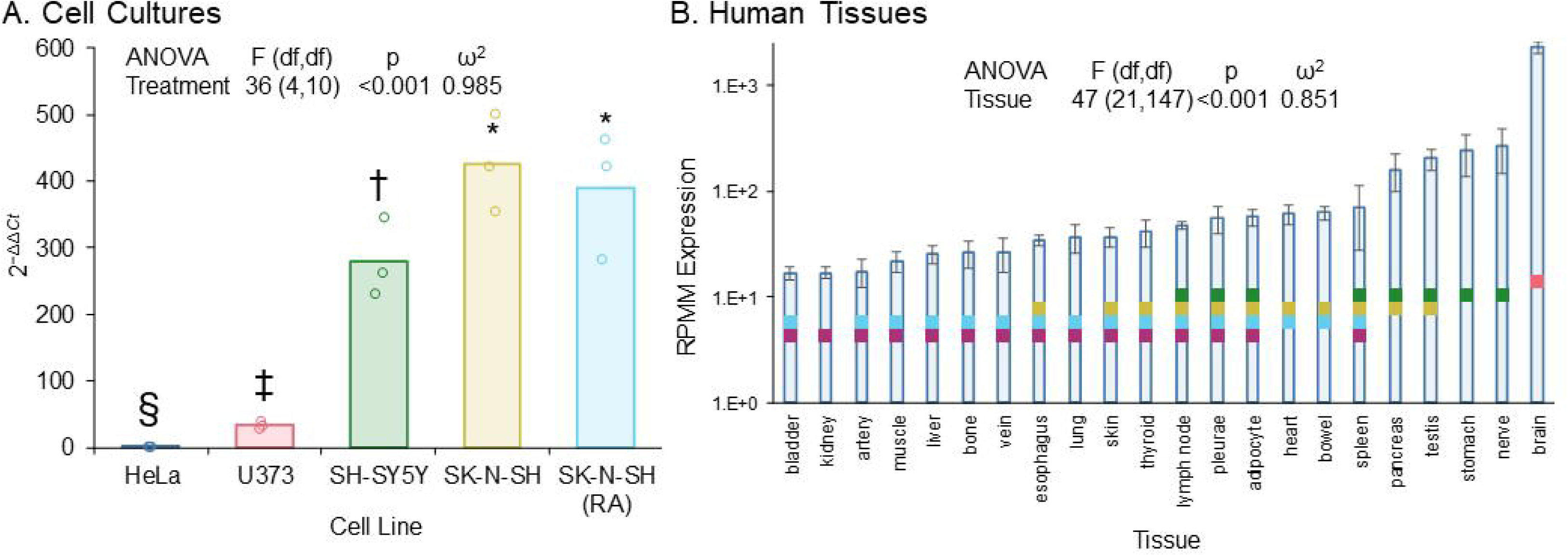
Levels of native miR-153-3p in various cell cultures and human tissues. We grew HeLa (epithelial), U373 (astrocytic), SK-N-SH, SK-N-SH stimulated with retinoic acid (RA), and SH-SY5Y (all three neuronal) cell lines, purified RNA, and measured levels of miR-153-3p by qRT-PCR. A) Neuronal cells had significantly higher levels of miR-153-3p than did non-neuronal. Symbols indicate differing statistical groups at p ≤ 0.05. Samples sharing symbols did not significantly differ. B) We queried the miRNATissueAtlas2 database for levels of miR-153-3p in human tissues. Modeling by tissue revealed significantly higher levels of the miRNA in brain vs. all other tissues.

### Levels of native miR-153-3p differ by tissue type

We compared levels of miR-153-3p by tissue type using glm, followed by all-vs-all pairwise comparisons. Our analysis revealed that brain had the highest level of miR-153-3p, followed by peripheral nerves and the stomach (Fig. 10B). Pairwise comparisons had poor resolution except that levels in brain were significantly higher than in all other tissues.

## Discussion

AD is a progressive degenerative disease resulting in neuron loss and synaptic disconnection. In this regard, recent studies have focused on mechanisms of neurogenesis, neural stem cell proliferation, and neuronal differentiation that might be harnessed in the AD brain to maintain neural circuits and stabilize cognitive deterioration [101, 102]. It was demonstrated that adult hippocampal neurogenesis is significantly reduced in AD patients as compared to healthy aged individuals, and that the number of doublecortin positive neural stem cells were significantly lower in the hippocampus of AD patients [103]. Doublecortin is required for normal migration of neurons into the cerebral cortex, since mutations in the human gene cause a disruption of cortical neuronal migration. The number of doublecortin and Proliferating cell nuclear antigen (PCNA) positive neurons was reduced in subjects with MCI and were associated with poorer cognitive function [104]. These data suggest the inhibition of neurogenesis from neural stem cells could contribute to AD progression and cognitive decline and, hence, restoring neurogenesis and increasing neural stem cell proliferation and differentiation could serve as a potential therapeutic strategy.

In the present work, we found that miR-153-3p regulates the expression of the transcription factor REST, which plays a critical role in neuronal maturation and maintenance, as well as several additional important proteins including SNCA and APP. Notably, human brain miR-153-3p levels also associated with decreased probability of AD. Furthermore, we found several AD-associated endophenotypes that were influenced by SNPs in proximity to the *MIR153-1* or *MIR153-2* genes, which were modified by the presence of *APOE*ε4 allele and biological variables such as age and sex.

To follow up on possible direct interactions between *APOE* and miR-153-3p, we explored *ad hoc* whether miR-153-3p directly interacted with the APOE mRNA sequence. Multiple bioinformatic prediction tools indicated that miR-153-3p has 2 predicted seedless binding sites on the APOE mRNA 3’-UTR, 1 seed-binding site on its CDS, 7 seedless binding sites on the CDS, as well as 1 seedless binding site on the 5’-UTR. On the other hand, the interactions we noted may reflect activity revealed when it was discovered that astrocytic ApoE protein delivers a variety of miRNAs into neurons, altering mRNA processing [105]. In particular, the *APOE* ε4-associated protein product has reduced capacity to deliver miRNAs into neurons compared to ε3 proteins. These potential multidirectional relationships warrant further investigation. We must note that our SNP analysis should not be considered exhaustive. We intentionally only highlighted the least entropically unfavored models and did not explore possible interactions of multiple SNPs with each other. Such a future comprehensive project would require even larger sample sets and computational power. Nevertheless, our findings could point to possibly screening SNP effects on miR-153-3p levels as the functional relationship between SNP and miR-153-3p disease effects.

In contrast to APOE, we located putative target sites in the mRNA 3’-UTR regions of REST, APP, and SNCA, and reporter clones with these UTRs responded in reporter assays to miR-153-3p treatment. In addition, disruptions of one site in the REST 3’-UTR reduced the effects of miR-153-3p on REST expression. Treatment of cell lines with miR-153-3p reduced levels of these proteins and associated native mRNA transcripts. These effects were seen in several cell lines of different tissue types. Furthermore, KEGG pathway analysis showed that treatment of differentiated neuroblastoma cells with miR-153-3p resulted in an enrichment of differentially expressed genes associated with axonal guidance, cell cycle, and focal adhesion. Notably, mutations in APP that alter its processing by secretases also alter activity of focal adhesion kinase, and focal adhesion regulates Aβ signaling and cell death in AD [106].Native levels of miR-153-3p indicated that the miRNA and its ensuing effects are not likely to be evenly distributed in all tissues. Our own comparison among epithelial, astrocytic, and neuroblastoma cells suggests enrichment in neuroblastoma lines, as we have found for other miRNA species, e.g., miR298 [107]. This was paralleled when we looked at levels of miR-153-3p in various tissues. Specifically, levels in brain were significantly and greatly elevated vs. peripheral tissues.

In neural stem cells, miR-153-3p treatment reduced nestin but increased doublecortin levels. REST knockdown by its siRNA reduced nestin but did not alter doublecortin levels. This indicated that miR-153-3p may regulate neuronal differentiation by both REST-dependent and independent pathways, which adds another layer of complexity that can be explored in future studies. For example, miR-153-3p induces differentiation of mouse hippocampal neuronal cells [108]. Bioinformatics indicates that miR-153-3p induces neuronal differentiation via promoting cell adhesion and BDNF/NTRK2 signaling pathway, while inhibiting ion channel activities in the cells [109]. miR-153-3p also promotes neurogenesis via targeting the notch signaling pathway. Inhibition of miR-153-3p reduces neural stem cell neurogenesis in hippocampus and leads to cognitive impairments [110]. In addition, miR-153-3p reduces nestin protein via targeting GPR55, a novel cannabinoid receptor.

REST not only can directly repress the expression of many neuronal genes but can also work together with miR124 and miR9 to fine-tune target gene expression levels [111–114]. REST represses miR124 expression, which reduces the expression of non-neuronal genes and induces neuronal mRNA profiling in HeLa cells [112]. REST also inhibits miR-9 transcription via binding to its promoter region. Reciprocally, miR9 binds to REST 3’ UTR and reduces REST expression [114]. Hence, miR-153-3p may act as a critical upstream regulator of neuronal integrity through multiple downstream pathways involving its regulation of REST.

With respect to study limitations, we recognize that several microRNAs play significant roles in the pathogenesis of AD [36–39, 41–45]. Interestingly, miRNAs fine-tune gene expression in different cell types via mRNA UTRs that are themselves subject to deletion or truncation [107]. We also recognize the common perception that 1) numerous miRNAs could potentially affect mRNA targets of miR-153-3p and, conversely, that 2) miR-153-3p regulates several targets. We address these points using the example of miR-153-3p and its target molecule APP. To test whether a target mRNA/protein is primarily regulated by miR-153-3p vs. several miRNAs, we have implemented our rigorous multi-step characterization protocols that counter unsupported claims of “a miRNA binds numerous targets.” For instance, it is true that bioinformatic prediction tools (e.g., TargetScan, miRanda, miRbase) with different algorithms predict several miR binding sites on target APP or even REST mRNA 3’UTRs. To address this specificity issue, we have used specific mRNA 3’-UTR-dual reporter systems to assay the functional activity of a particular 3’-UTR (or 5’-UTR [37]). Indeed, when we separately transfected several predicted miRNAs in neuronal cultures, many miRNAs did not regulate APP, underscoring our well-controlled mechanistic approach. Moreover, miR:mRNA:protein interactions and functional output are complex; UTR activity does not always mirror protein level. Therefore, we measure target protein levels upon detecting positive signals from UTR activity. The advantage of our strategy of studying miRNA effects on levels of native protein are different from typical studies using only exogenous UTR-reporter activity [49, 115]. The native protein level is regulated by endogenous miR levels and by those supplied exogenously. To test whether APP protein is regulated primarily by miR-153-3p, we transfected miR-153-3p mimic and observed both APP mRNA and native APP levels were significantly reduced. Notably, when anti-miR-153-3p was added, APP protein levels were almost restored to the mock level. Had several miRNAs regulated APP protein, then neutralizing one particular miR (in this case miR-153-3p) would not entirely reverse inhibition. Furthermore, we previously showed that miR-153-3p reduced levels of APP in neuron-enriched human primary mixed brain cultures, and here we report miR-153-3p’s effects in hiPSC and autopsied human brain tissue specimens [36].

An additional caveat is that there were no animal models used in this work. Unfortunately, conservation between human and mouse miR-153-3p binding sites on our target mRNAs 3’UTR can be low. Such low conservation reduces applicability of a naïve use of mouse models and complicates translation in a human setting. Finally, it is already established that miR-153-3p is extensively involved in a variety of cancers and targets the expression of multiple oncogenes, acting as an oncogenic or tumor suppressing miRNA depending on the type of cancer, where it can differentially regulate tumor proliferation, invasion, angiogenesis, apoptosis and autophagy [116]. On the other hand, sponging of miR-153-3p by non-coding RNAs modulates tumorigenicity activity [116]. Thus, the activity on miRNA may be another factor in the inverse risks of cancer and AD [117, 118].

In summary, we have demonstrated that miR-153-3p is such a multi-targeting miRNA has activity in reducing REST, SNCA, and APP levels via 3’UTR-directed regulation of their respective mRNAs and that increased miR-153-3p levels in cortex are associated with a decreased probability of AD. Moreover, we identified SNPs within 5kb of the *MIR153-1* and *MIR153-2* genes that were significantly associated with changes in AD-associated endophenotypes. Finally, we provided evidence that miR-153-3p regulates the neuronal phenotype via reducing REST, increasing doublecortin expression, and inducing genes associated with axonal guidance, cell cycle, and focal adhesion. Our work suggests that miR-153-3p supplementation will reduce levels of toxic protein aggregates by decreasing the expression of APP and SNCA and promote neuronal maintenance and survival by reducing REST.

The current findings underscore the progress we have made in discovering several miRNAs that specifically regulate translation of important genes associated with AD (e.g., APP, BACE1, MAPT, and REST). Altogether, mechanistic studies in cell cultures, manipulations in iPSCs, miRNA expression patterns in AD brains, and SNPs identification in MIRNA genes will establish their relationships to AD pathogenesis. Uniting our own work with that known to the field, we propose a mechanistic cascade to explain the roles of miR-153-3p in AD (Fig. 11). In this model, miR-153-3p would serve to target the RISC to (among others) REST, APP, and SNCA mRNA 3’-UTR sequences. This blockade would reduce levels of APP and resultant Aβ and levels of SNCA. Blockade of REST would influence neuronal differentiation and cell health. Overall REST transcription regulation may, itself, influence miR-153-3p production. Dysregulation of miR-153-3p would perturb normal regulation, resulting in CNS dysfunction. Taken together, these data point towards the therapeutic and biomarker potential of targeting miR-153-3p in AD and related dementias.

**Fig. 11.**
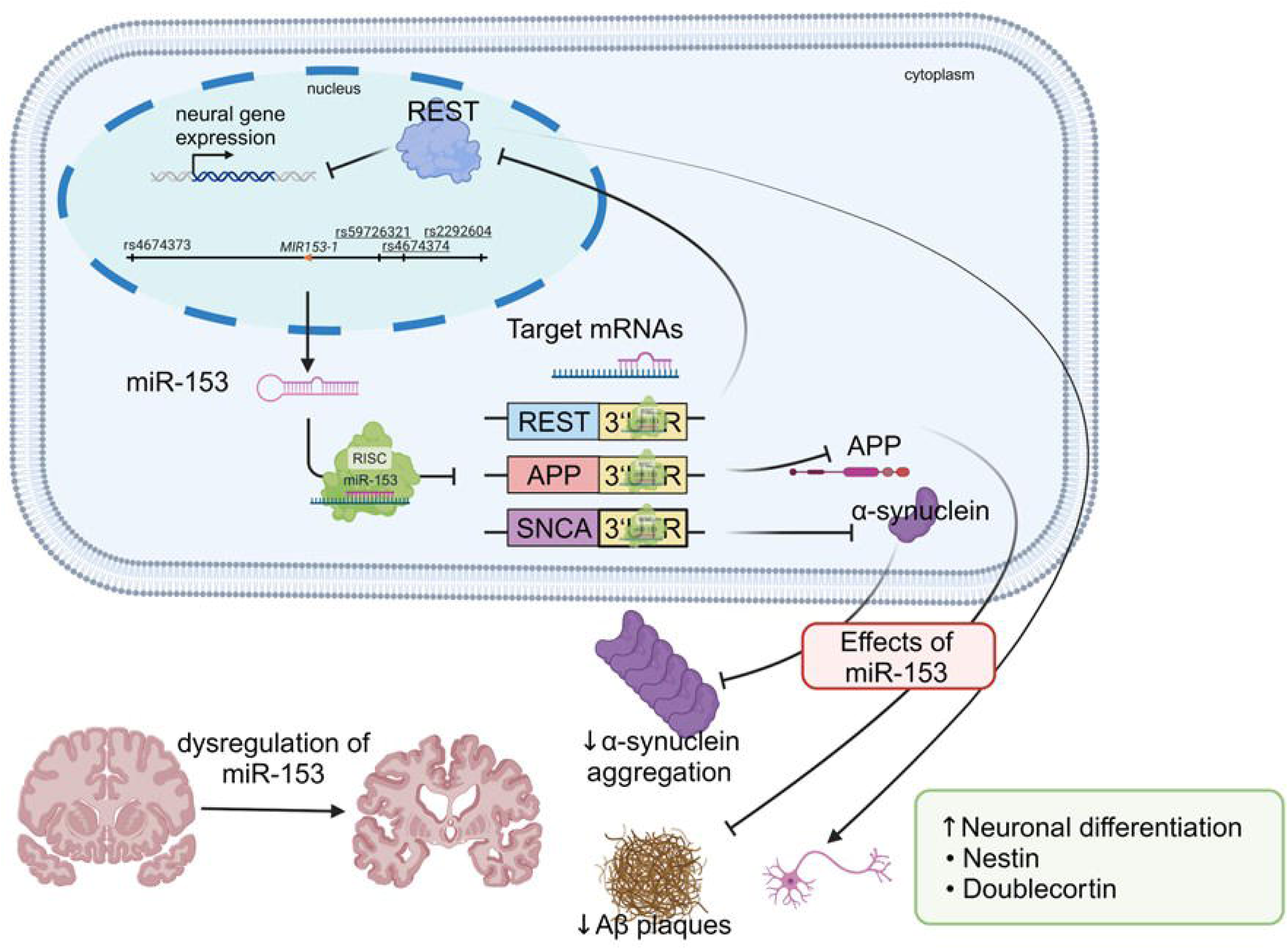
Roles of miR-153-3p in connection to neurodegeneration and AD. Based on our own work and others known to the field, we propose a mechanistic cascade to explain the roles of miR-153-3p in AD. In this model, miR-153-3p would serve to target the RISC to (among others) REST, APP, and SNCA mRNA 3’-UTR sequences. This blockade would reduce levels of APP and resultant Aβ and levels of SNCA. Blockade of REST would influence neuronal differentiation and cell health. Overall REST transcription regulation may, itself, influence miR-153-3p production. Dysregulation of miR-153-3p would perturb normal regulation, resulting in CNS dysfunction.

In addition to microRNAs regulating mRNA translation, natural antisense transcripts (NATs) regulate mRNA levels, and translation. Furthermore, NATs are noncoding RNAs (ncRNAs) encoded by DNA sequences overlapping the pertinent protein genes. NAT’s roles in regulating key protein-coding genes associated with neurodegenerative disorders are emerging [119]. Future research in elucidating the functions of NATs and miRNAs could prove useful in treating or preventing diseases.

## Acknowledgements

Sincere thanks to Drs. Bernardino Ghetti, Martin Farlow, Peter Nelson, Andrew Saykin, and Alzheimer’s Disease Neuroimaging Initiative (ADNI). Some of the data used in the preparation of this article were obtained from the NIH-funded ADNI database (adni.loni.usc.edu). For up-to-date information, see adni.loni.usc.edu.

